# Mouse models of COVID-19 recapitulate inflammatory pathways rather than gene expression

**DOI:** 10.1101/2022.02.24.481866

**Authors:** Cameron R. Bishop, Troy Dumenil, Daniel J. Rawle, Thuy T. Le, Kexin Yan, Bing Tang, Gunter Hartel, Andreas Suhrbier

**Affiliations:** Immunology Department, QIMR Berghofer Medical Research Institute, Brisbane, Queensland. 4029, Australia; Australian Infectious Disease Research Centre, GVN Center of Excellence, Brisbane, Queensland, 4029 and 4072, Australia

## Abstract

**BACKGROUND:** How well mouse models recapitulate the transcriptional profiles seen in humans remains debatable, with both conservation and diversity identified in various settings. The K18-hACE2 mouse model has been widely used for evaluation of new interventions for COVID-19.

**METHOD:** Herein we use RNA-Seq data and bioinformatics approaches to compare the transcriptional responses in the SARS-CoV-2 infected lungs of K18-hACE2 mice with those seen in humans.

**RESULTS:** Overlap in differentially expressed genes was generally poor (≈20-30%), even when multiple studies were combined. The overlap was not substantially improved when a second mouse model was examined wherein hACE was expressed from the mouse ACE2 promoter. In contrast, analyses of immune signatures and inflammatory pathways illustrated highly significant concordances between the species.

**CONCLUSION:** As immunity and immunopathology are the focus of most studies, these hACE2 transgenic mouse models can thus be viewed as representative and relevant models of COVID-19.

## Introduction

Mouse models represent critical tools for preclinical evaluation of new interventions and for understanding *inter alia* disease, host responses and pathogen behaviors. However, views on how well mice recapitulate human transcriptional profiles range from substantial conservation (1, 2) to considerable diversity (3). In pro-inflammatory settings, reports have also argued that mouse models mimic human transcriptional responses well (4) or poorly (5). Mouse models of human disease can thus be seen as less reliable (6–10) or, in other settings, as recapitulating faithfully key elements of human disease (11–13). Given both cross species conservation and diversity can be identified (14, 15), specifically interrogating any given mouse model for how reliably its transcriptomic responses mimic those seen in humans is clearly warranted (16–18).

A widely used mouse model of SARS-CoV-2 infection and COVID-19 disease is the K18-hACE2 mouse, where the human angiotensin-converting enzyme 2 (hACE2) is expressed as a transgene from the keratin 18 (K18) promoter. These mice develop a robust respiratory disease that histologically resembles severe COVID-19 (19–21). These mice have been widely used for evaluation of new interventions (22–27), and for virology and immunopathology studies (28, 29). However, SARS-CoV-2 infected K18-hACE2 mice show a number of differences (19), perhaps the most important difference is a fulminant brain infection that is associated with the generally lethal outcome in this model (30). Although human brain infection has now been demonstrated (31), fulminant lethal brain infection is not a feature of COVID-19 (32, 33). A second mouse model of SARS-CoV-2 infection and COVID-19 disease involves expression (also as a transgene) of hACE2 driven by the mouse ACE2 promoter (34), referred to herein as mACE2-hACE2 mice. This mouse model does not show the brain infection seen in K18-hACE2 mice, with infection generally self-limiting and non-lethal (34). Herein we describe a bioinformatic approach for comparing the transcriptional responses of mouse lungs and human lungs/lung tissues after SARS-CoV-2 infection. Although overlap for differentially expressed genes (DEGs) across species was generally poor, concordance for immune signatures and inflammation pathways was high.

## Results

### Mouse and human data sets for SARS-CoV-2 infected lung tissues

RNA-Seq datasets for lungs from SARS-CoV-2 infected K18-hACE2 mice were obtained from two independent sources, Winkler and our own group, Suhrbier (Table 1). For the former (35), datasets for lungs of SARS-CoV-2 infected K18-hACE2 mice for 2, 4 and 7 days post infection (dpi), were obtained from the NCBI Sequence Read Archive (SRA) (Table 1). Two Suhrbier K18-hACE2 datasets were available, 2 and 4 dpi, with the latter reported previously (27) and the former generated for this study (Table 1). Fastq files were analyzed or reanalyzed herein using STAR, RSEM and EdgeR, with a q<0.05 filter applied to provide Differentially Expressed Genes (DEGs) (Supplemental Table 1 a, d, g, j, m). For each of these DEG lists, a mouse-human orthologue DEG list (orthoDEGs) (Supplemental Table 1 b, e, h, k, n) and a single copy orthologue DEG list (scoDEGs) was generated (Supplemental Table 1 c, f, i, l, o).

**Table 1:** Origins of human and K18-hACE2 gene expression datasets. K18-hACE2 mouse studies from two independent laboratories provided five data sets from lungs of SARS-CoV-2-infected K18-hACE2 mice. Mouse data covered 3 time points representing days post infection (dpi). Four independent human studies provided three gene expression datasets from SARS-CoV-2-infected lungs, and one from 3D lung organ culture infected *ex vivo* with SARS-CoV-2. The Ackerman study analysed expression of 249 mRNAs associated with inflammation. All infections were with SARS-CoV-2 isolates belonging to the original or ancestral lineage. PRJ prefixed annotations represent NCBI Bioproject accession numbers. Reanalysis means raw fastq files were re-analyzed for this study using STAR, RSEM and EdgeR.

| Study name | Species | Dataset source | Lung Tissues | Method | Platform | Re-analysed | Notes |
| --- | --- | --- | --- | --- | --- | --- | --- |
| Winkler 2, 4 & 7 dpi | Mouse (K18-hACE2) | PRJNA645133 | SARS-CoV-2 vs mock infected lung | RNA-Seq (poly-A selected) | Illumina NovaSeq 6000 | Y | 2.5x10<sup>4</sup> p.f.u. n=4-6 per group |
| Suhrbier 2 & 4 dpi | Mouse (K18-hACE2) | PRJNA767499 PRJEB43658 | SARS-CoV-2 vs mock infected lung | RNA-Seq (poly-A selected) | Illumina NextSeq 500 | Y | 5x10<sup>4</sup> CCID<sub>50</sub> n=4-6 per group |
| mACE2-hACE2 2, 4, 6 & 10 dpi | Mouse (mACE2-hACE2) | PRJNA767499 | SARS-CoV-2 vs mock infected lung | RNA-Seq (poly-A selected) | Illumina NovaSeq 6000 | Y | 5x10<sup>4</sup> CCID<sub>50</sub> n=3/4 per group |
| Wu | Human | PRJNA646224 | Formalin fixed paraffin embedded post mortem COVID-19 vs control biopsies | RNA-Seq (rRNA-depleted) | Illumina NextSeq 550 | Y | Infected n=9, Control n=10. Medicated. |
| Alfi | Human | PRJNA688321 | *Ex vivo* 3D lung organ cultures, infected vs mock | RNA-Seq (poly-A selected) | Illumina NextSeq 500 | Y | Infected n=10 Control n=10 |
| Blanco-Melo | Human | PRJNA615032 | Formalin fixed post mortem COVID-19 vs control lung | RNA-Seq (poly-A selected) | Illumina NextSeq 500 | N | Infected n=2 Control n=2 4 RNA-Seq libraries for each. |
| Ackermann | Human | Vivli Center for Clinical Research Data | Formalin fixed post mortem COVID-19 vs control lung | Nano-String | nCounter® Inflammation Panel | N | Infected n=7, Control n=10 Alveolar damage noted |

Three RNA-Seq datasets derived from SARS-CoV-2 infected human lung samples (COVID-19 vs. Controls) were obtained from the NCBI Sequence Read Archive (SRA) and are referred to as Wu (36), Alfi (37) and Blanco-Melo (38) (Table 1). The Alfi and Wu data sets were reanalyzed (using STAR, RSEM and EdgeR) to produce gene expression datasets. Compared to the Alfi and Wu data sets, the Blanco-Melo dataset showed very low sequencing depth in the COVID-19 samples (Supplemental Figure 1), and the original gene list provided by the authors (38) was used. DEGs (Supplemental Table 1p, s, v), orthoDEGs (Supplemental Table 1q, t, w) and scoDEGs (Supplemental Table 1r, u, x) were generated as above for the 3 human groups, respectively. An additional human COVID-19 DEG list derived from NanoString analysis (39) was obtained from the Vivli Center for Clinical Research Data (Table 1, Ackermann).

In summary, five DEG, orthoDEG and scoDEG datasets for 3 time points and from two independent groups, describe significant differential gene expression in lungs of SARS-CoV-2-infected K18-hACE2 mice. Four independent DEG, orthoDEG and scoDEG datasets were obtained that describe significant differential gene expression in SARS-CoV-2-infected human lung tissues. The number of DEGs ranged from 75 to 2794 per dataset (Figure 1A). The proportion of DEGs in each dataset that were orthoDEGs or scoDEGs between mouse and human ranged from 57 to 92% and 49 to 83%, respectively (Figure 1A). When the NanoString (Ackerman) and the Organ culture (Alfi) was removed, these percentages were 74 to 83% and 63 to 77%, respectively (Figure 1A).

**Figure 1:**
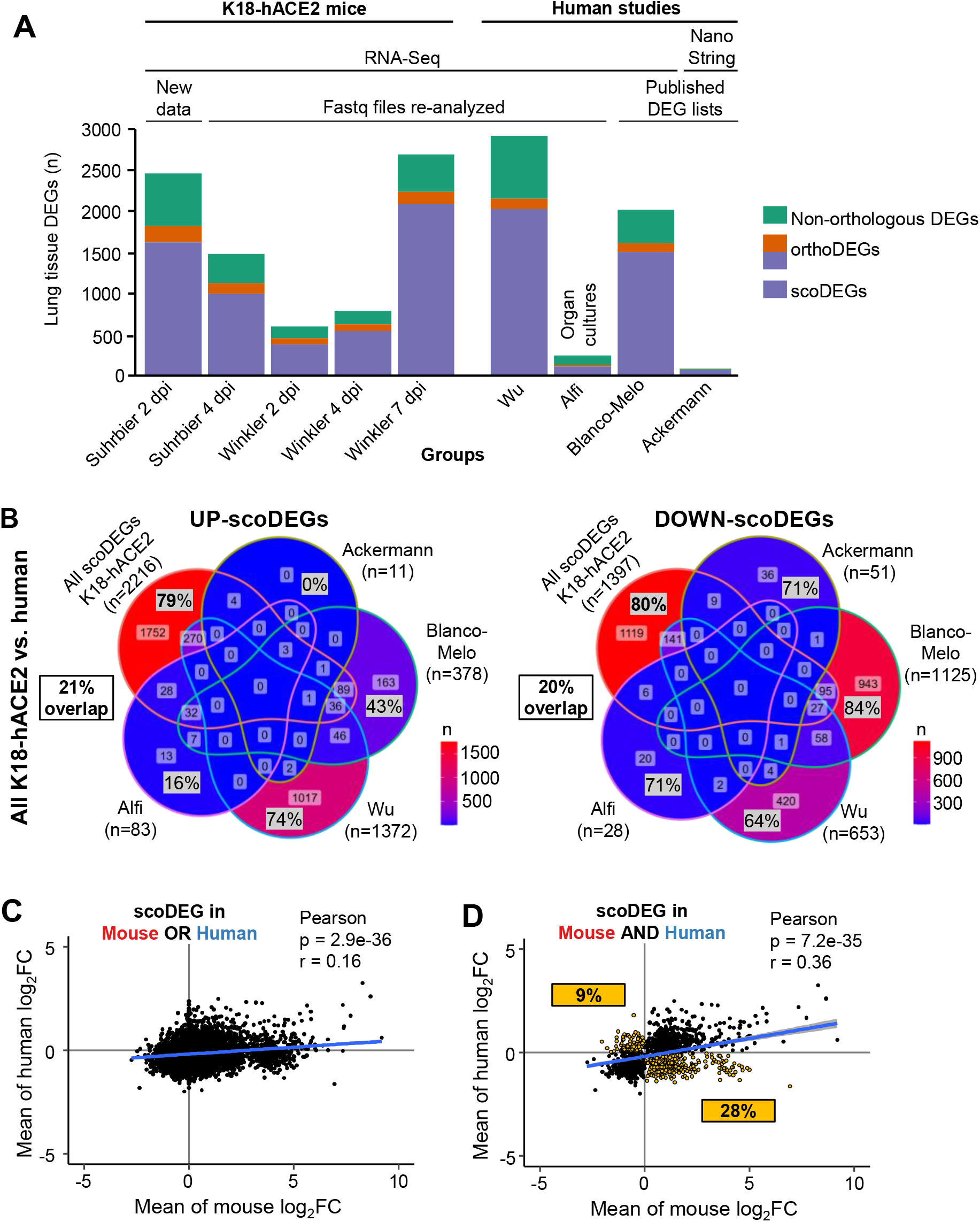
Mouse and human expression data sources and degree of concordance between species. **(A)** DEGs from lungs/lung tissues infected with SARS-CoV-2 were identified in K18-hACE2 mouse and human studies (n= number of DEGs). DEGs were generated from original RNA-Seq data provided herein (New Data), re-analyzed from previously published RNA-Seq data (Fastq files re-analysed), or were obtained from publications (published DEG lists). All but one dataset were derived from RNA-Seq and one was derived from a microarray study. Green - non-orthologous between mouse and human. Orange - with one or both species having multiple orthologues. Purple - both species having a single copy orthologue. A total of 9 Groups (5 K18-hACE2 and 4 human) were considered in the subsequent analyses. **(B)** The union of all K18-hACE2 scoDEGs was used to compare mouse and human for up- and down-regulated scoDEGs. ‘n’ refers to the number of scoDEGs for each group. Percentages within the Venn diagram (gray boxes) show the percentage of scoDEGs exclusive to that group (i.e. a scoDEG in that group but no other group) (e.g. 1752/2216 × 100 = 79%). The boxed overlap percentages represent the percentage of mouse scoDEGs that are also scoDEGs in one or more human studies (e.g. 2216-1752/2216 × 100 ≈ 21% for up-regulated scoDEGS and 1119-1397/1397 ×100 ≈20% for down-regulated scoDEGs). **(C)** Pearson correlation of mean log_2_FC changes of single-copy orthologues that were DEGs in either any mouse group or any human group or both. **(D)** Pearson correlation of mean log_2_FCs of single-copy orthologues that were DEGs in both one or more mouse groups and one or more human groups. ScoDEGs that had inconsistent mean expression between species (i.e. were upregulated in one species and down-regulated in another) are shown yellow. The percentage of scoDEGs with inconsistent expression (yellow boxes) is provided relative to the total number of scoDEGs.

### Poor overlap between human and mouse DEGs

When the up-regulated scoDEGs from all the mouse data sets were combined (n=2216) and compared with scoDEGs from each of the four human studies, 79% of scoDEGs up-regulated in mice were not up-regulated in any human study (Figure 1B). The same comparison for down-regulated scoDEGs showed 80% of scoDEGs down-regulated in mice were not down-regulated in any human study (Figure 1B). Thus the overall overlap for up- and down-regulated scoDEGs for K18-hACE2 mice and human studies was only 21 and 20%, respectively.

Conceivably, due to the q<0.05 cutoff, a DEG in one species may narrowly have missed being a DEG in the other species because it just missed out on significance. A K18-hACE2 mouse-vs. human comparison was thus undertaken using single-copy orthologues that were differentially expressed in at least one mouse group OR at least one human group (union scoDEGs) (Figure 1C), and a second comparison using single-copy orthologues that were differentially expressed in at least one mouse group AND at least one human group (intersection scoDEGs) (Figure 1D). For each comparison, log_2_ fold-changes (log_2_FC) were averaged within each species (Supplemental Table 2) and tested for correlation. Although significant, correlation coefficients were relatively poor in both comparisons; r=0.16 and 0.36 for union scoDEGs and intersection scoDEGs, respectively (Fig 1C, D). The poor overlap between DEGs in K18-hACE2 mice and human studies cannot therefore be readily explained by genes just missing the p<0.05 cutoff. In addition, 37% (9% + 28%) of genes showed opposite directions of average fold change for mouse and human scoDEGs (Figure 1D, yellow boxes).

When DEG overlaps were calculated in pairwise comparisons between each group, overlaps between mouse and human data sets remained low, ranging from 1-9% (Supplemental Figure 2). This illustrated that no human group showed good concordance with any mouse group, and that overlaps were higher when multiple studies were combined (Figure 1B). Overall these analyses argue that differential gene expression in SARS-CoV-2 infected K18-hACE2 lungs, by enlarge, do not recapitulate particularly well differential gene expression in lungs of infected human lung tissues.

### Large variations in viral read counts

To assess the viral loads for each group, the percentage of reads mapping to the virus was determined for each RNA-Seq dataset and expressed as a percentage of reads aligned to protein-coding genes. Mouse groups had ≈1.5 - 5.5 logs more mean viral read counts than the human studies (Figure 2A). The high viral read counts in K18-hACE2 mice is perhaps not surprising given that this is a robust lethal model of SARS-CoV-2 infection (20, 28, 35).

**Figure 2:**
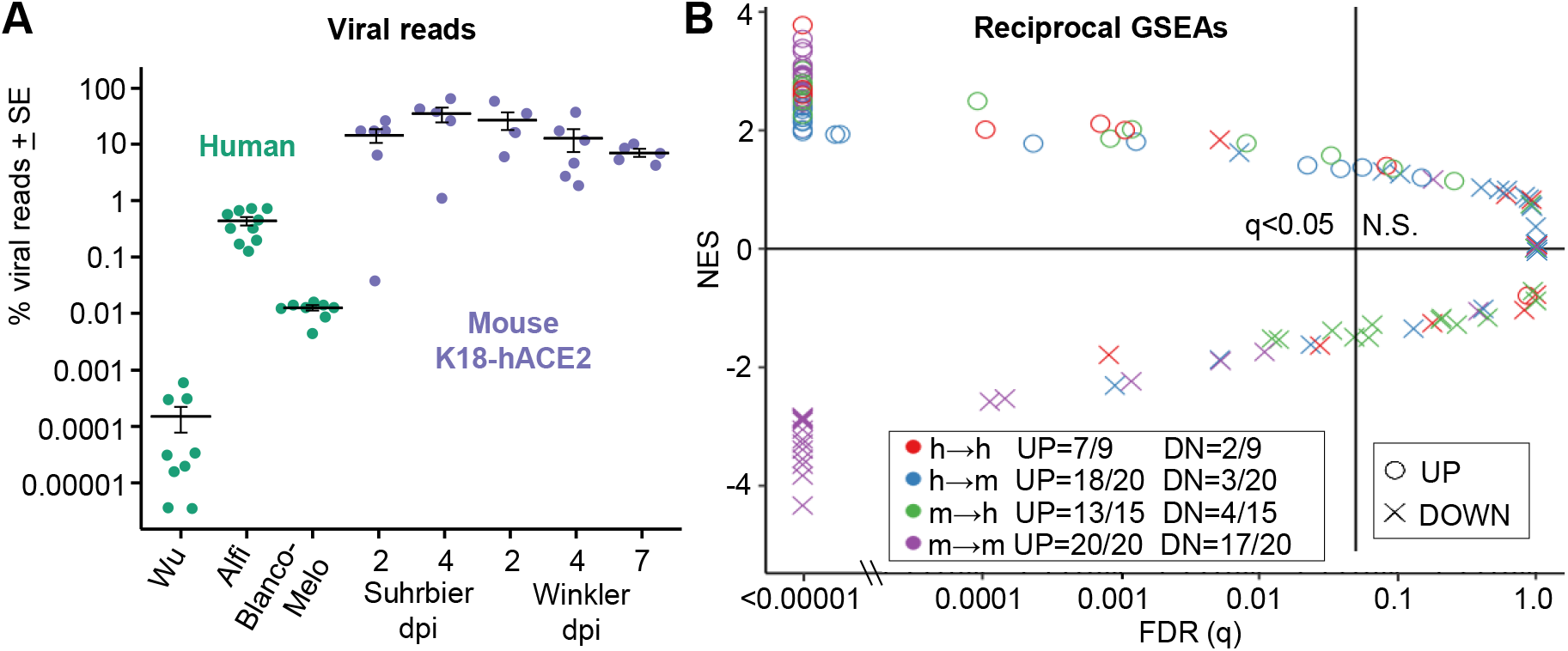
Viral reads and reciprocal GSEAs. **(A)** For each sample, the number of reads aligned to the SARS-CoV-2 genome are shown as a percentage of the total number of reads that align to all protein coding genes (filled circles). Cross-bars represent the mean for each group. **(B)** Pairwise reciprocal GSEAs showing enrichment of up- or down-regulated orthoDEG sets in log_2_FC ranked gene lists for all possible pair-wise comparisons between groups (i.e. 128 combinations; 3 human and 5 mouse ranked gene lists vs. 4 human and 5 mouse orthoDEG lists). Circles and crosses are colored according to direction of the GSEA (e.g. green = mouse orthoDEG sets vs. ranked human gene lists). Fractions show the proportion of orthoDEG sets that are significantly enriched with consistent directionality (e.g. “UP=13/15” indicates that, of 15 GSEAs using up-regulated orthoDEG sets, 13 showed significant enrichment with positive NES).

The analysis also illustrated the large differences in viral loads between human studies, Alfi contained ≈2 logs more viral reads than Blanco-Melo, and Blanco-Melo had ≈2 logs more viral reads than Wu. In contrast, the mean percent read counts varied by only ≈1 log between mouse groups (Figure 2A). The large differences in viral loads in the human groups may explain, at least in part, why DEG overlap between any of the human groups was low (Supplemental Figure 3A). The lower differences in viral loads in the mouse groups may explain, at least in part, why DEG overlap between any of the mouse groups was higher (Supplemental Figure 3B). However, as illustrated below, the poor scoDEG overlap between K18-hACE2 and human groups was not substantially rectified when the non-lethal mACE2-hACE2 model was used, which has significantly lower lung titers than K18-ACE2 mice.

### Gene Set Enrichment Analyses using orthoDEGs

A method for comparing gene expression data sets is to use Gene Set Enrichment Analysis (GSEA), whereby the enrichment of a DEG set within a pre-ranked gene list is evaluated (40–43). For orthologues with different gene nomenclature in mice and humans, the mouse gene symbols were changed to their orthologous human equivalent in the orthoDEG sets and the gene lists. This allows GSEAs to be undertaken for mouse vs. human gene sets. The orthoDEG gene sets used in the GSEAs comprised the top 50% of orthoDEGs ranked by fold change (Supplemental Table 1). These orthoDEG sets were then used to interrogate all other ranked gene lists (ranked by fold change) for all the other groups (Supplemental Table 3).

The up-regulated orthoDEG sets from almost all groups were significantly enriched with positive normalized enrichment scores (NES) in all the ranked gene lists (Figure 2B, circles; Supplemental Table 4). Of the 35 (20 plus 15) GSEAs for mouse-human and human-mouse comparisons, 31 (18 plus 13) reached significance (Figure 2B, q<0.05 blue and green circles). Thus although overlap in up-regulated orthoDEGs was poor and variation in viral loads was high, the top up-regulated orthoDEGs identified in SARS-CoV-2 infected lungs from K18-hACE2 mice generally showed significant enrichment in human ranked gene lists and *vice versa*.

GSEAs using down-regulated orthoDEGs provided the opposite result for mouse-human and human-mouse comparisons with only 7 (3 plus 4) out of 35 GSEAs reaching significance (Figure 2B, q<0.05, blue and green crosses; Supplemental Table 4). In addition, only 2/9 human-human GSEAs of down-regulated orthoDEGs reached significance (Figure 2B, red crosses). However, for mouse-mouse GSEAs, the number that reached significance for down-regulated orthoDEGs (17/20) was only marginally lower than for up-regulated orthoDEGs (20/20) (Figure 2B).

Analyses of down-regulated DEGs by Ingenuity Pathway Analysis (IPA) Diseases and Functions feature, remarkably provided the same top scoring annotation by z-score for all groups, Organismal death, with the exception of the Alfi organoid study where the top annotation was Perinatal death (Supplemental Table 5). Down-regulated DEGs would thus largely and consistently appear to arise from tissue damage, with cell death and injury in SARS-CoV-2 infected lungs induced via direct viral cytopathic effects (44) and via the cytokine storm (45). The comparable levels of virus infection seen in the mouse groups (Fig. 2A) might thus contribute to the higher level of congruence (i.e. 17/20 significant GSEAs) in the down-regulated gene signatures (Figure 2B, purple crosses).

### GSEAs using ImmuneSigDB shows concordance between mouse and human studies

ImmuneSigDB is a compendium of ≈5000 immunology-specific gene sets that can be used to interrogate mouse and human ranked gene lists using GSEAs to identify immune signatures (15). GSEAs using ImmuneSigDB gene sets were used to interrogate the gene lists ranked by fold change (Supplemental Table 3). The NES are plotted for GSEAs that reached significance (q<0.05), with the NES ranked by Suhrbier 4 dpi (Figure 3A), as this dataset provided the largest number of significant GSEA results (n=1969). The results were also grouped by ImmuneSigDB gene sets that mentioned a specific cell type (Figure 3A) in the gene set annotation (15). Concordance between the mouse and human GSEA results was generally high, with the concordance also apparent across all cell types (Figure 3A).

**Figure 3.**
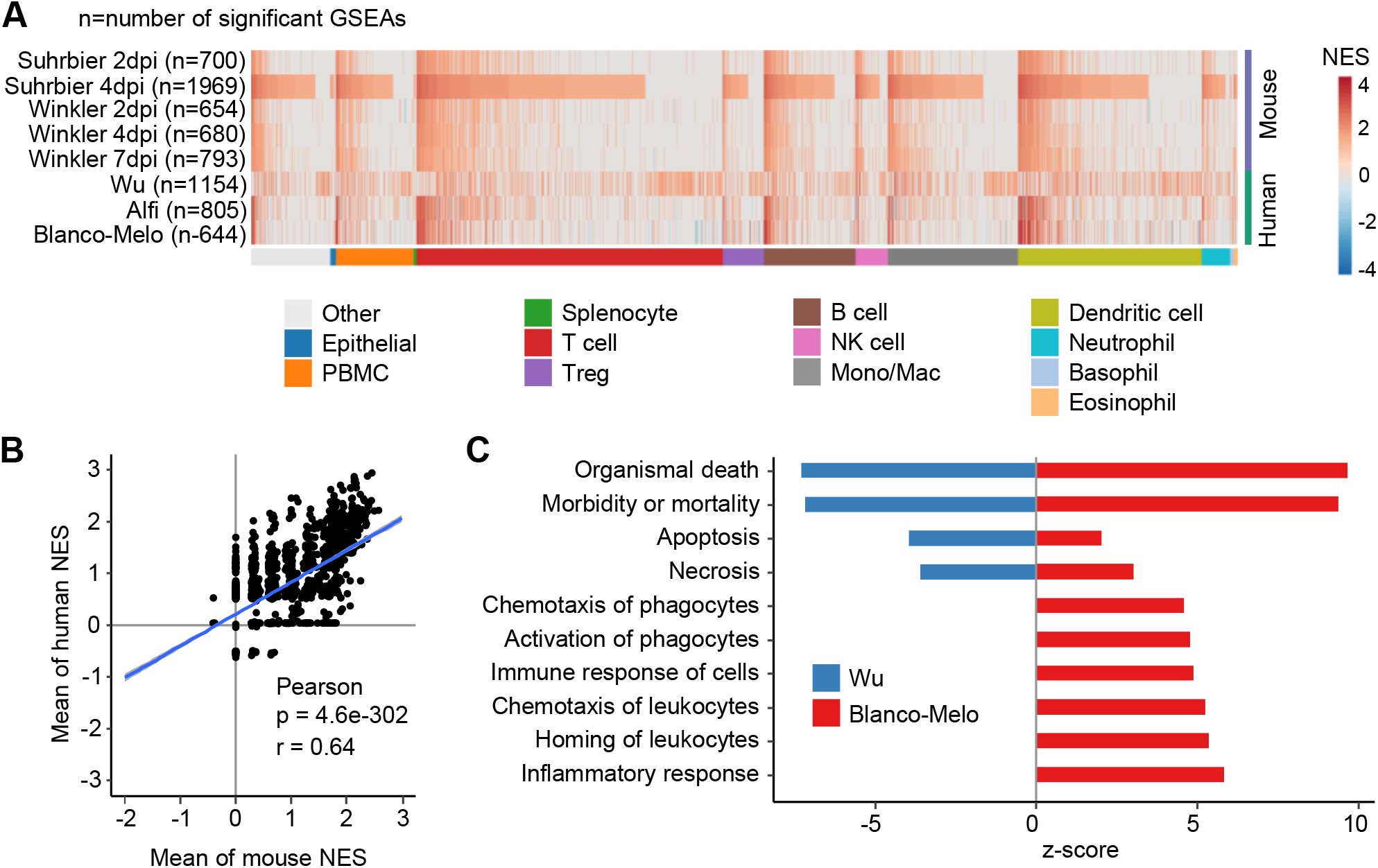
Gene Set Enrichment Analysis of immune-cell related gene sets. **(A)** Gene sets from the GSEA Immune Signatures Database (ImmuneSigDB) were used to interrogate log2FC ranked gene lists from all groups. A total of 2879 GSEAs were significantly enriched in at least one group, and were clustered according to cell-type. Within each cell-type cluster, Gene sets were ranked according NES for Suhrbier 4 dpi. **(B)** Pearson’s correlation of mean mouse NES for 2879 ImmuneSigDB gene sets vs. the mean human NES for the same gene sets. **(C)** The Wu and Blanco-Melo DEGs were analysed by IPA Diseases and Functions. The top 10 annotations that had the greatest differences in z-scores between Wu and Blanco-Melo are shown, ranked by difference. A z-score of 0 means the annotation was not identified as significant by IPA.

A Pearson correlation of the mean NES for human vs mouse provided a highly significant correlation (Figure 3B, r=0.64), illustrating that both human and mouse datasets showed comparable enrichments for many ImmuneSigDB gene sets. In summary, despite the poor overlap for scoDEGs, GSEAs using ImmuneSigDB gene sets argue that human and mouse ranked gene lists share a significant number of immune related signatures.

The Wu dataset showed a clearly distinct pattern (Figure 3A). To gain insights into why, the DEGs from the Wu and Blanco-Melo studies were analyzed by IPA Diseases and Functions. The results were ranked by difference in z-scores, with the top 10 most different annotations shown (Figure 3C). Cell death and inflammation annotations had much lower z-scores in the Wu study when compared with the Blanco-Melo study (Figure 3C). These results are consistent with the ≈2 log lower mean level of virus in the Wu group (Figure 2A), with methylprednisolone treatments in 4/9 patients (and interferon in one patient and intravenous immunoglobulin in another) (36) perhaps also suppressing inflammation and thereby prolonging survival (46) to a time when viral titers have waned. The antiviral treatments used in this study (36) have subsequently been shown not to have significant activity.

### Cytokine/chemokine signaling pathways show high level concordance between species

For COVID-19 a key feature is up-regulation of inflammatory mediators, with the ensuing cytokine storm associated with acute respiratory distress syndrome (ARDS) that characterizes severe disease (47). The 5 mouse and 4 human DEG sets were analyzed by IPA, which accepts both human and mouse gene nomenclature. The Up-Stream Regulator (USR) feature of IPA provided a list of z-scores for cytokine (which includes chemokine) pathways, which were then ranked and plotted in heat maps. This analysis illustrated considerable concordance between the dominant pro-inflammatory cytokine/chemokine responses in mice and humans (Figure 4A,B). A highly significant correlation emerged between the mean z-scores for cytokine/chemokine USRs for human groups and the mean z-score for cytokine/chemokine USRs for mouse groups (Figure 4C, Cytokine/chemokine). This correlation remained highly significant when cytokine/chemokine USR z-scores from individual mouse data sets were used instead of the means (Supplemental Figure 4). Thus, although orthoDEGs and scoDEGs showed poor overlaps for human and mouse groups, pathway analyses illustrated that the cytokine/chemokine responses in SARS-infected human and mouse lung tissues were actually quite similar.

**Figure 4:**
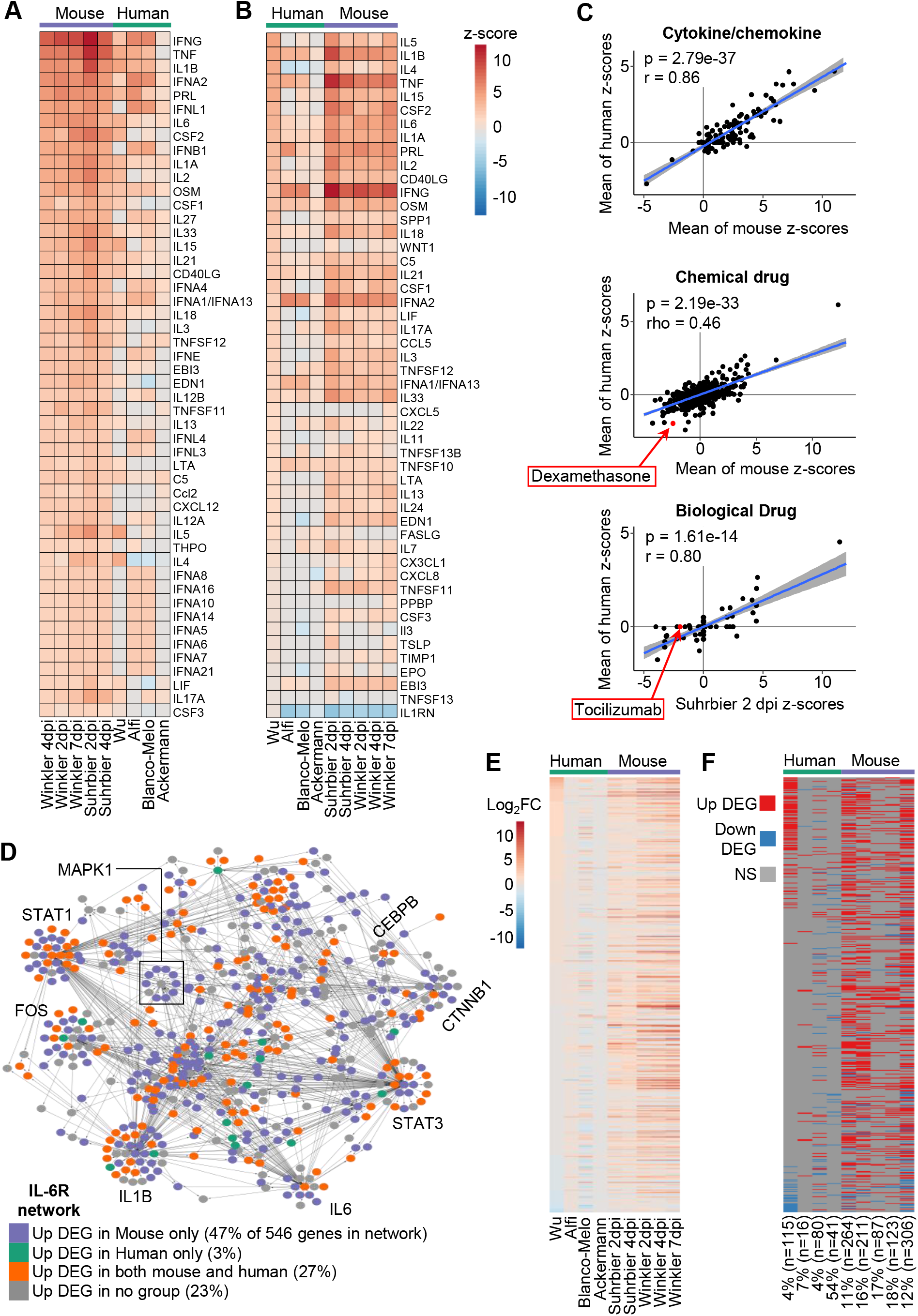
Cytokine/chemokine and drug USR concordances between humans and K18-hACE2 mice. **(A)** DEGs from each group were analysed by IPA upstream regulator (USR) feature. The heatmap shows the top 50 of cytokine/chemokine USRs ranked by activation z-scores from the Winkler 4 dpi data. **(B)** Heatmap comparing groups as in A, except ranked according to z-score from the Wu data. **(C)** Cytokine/chemokine – Pearson correlation of mean mouse z-scores vs. mean human z-scores for significant Cytokine/chemokine USRs (n=121). Chemical drugs – Spearman correlation of mean mouse z-scores vs. mean human z-scores for significant Chemical drug USRs (n=624). Biologic drugs - Pearson correlation of mean mouse z-scores vs. mean human z-scores for significant Cytokine/chemokine USRs (n=59). For calculating means, non-significant USRs were given a value of zero, thus means were derived from n=5 for mouse groups and n=4 for human groups. **(D)** Network of 546 genes associated with IL-6R signaling according to IPA. Node color indicates whether a gene was up-regulated in mouse only (purple; >1 mouse group, and no human group), human only (green; >1 human group, and no mouse group), both (orange; >1 mouse and >1 human group), or not up-regulated in any group (grey). Large sub-networks are labeled according to their hub node. **(E)** Heatmap comparing groups according to log_2_ fold-change (log_2_FC) of 546 genes associated with the IL-6R signaling network. Genes are ranked according to log_2_FC in Wu. **(F)** Categorical “heatmap” with genes ordered as in E, with up-regulated DEGs shown in red, down-regulated DEGs in blue, and genes whose expression was not significantly different (NS) in grey. The number of IL-6R network genes that were DEGs is shown for each group as a percentage of the total number of DEGs. The total number of DEGs is also provided (n).

A series of human clinical trials have shown the benefit of anti-inflammatory treatments for COVID-19 ARDS such as corticosteroids (e.g. dexamethasone) (48) and anti-IL-6-receptor (tocilizumab) (49). When IPA Chemical Drug USRs were compared, a highly significant correlation emerged, with dexamethasone appearing with the expected negative z-score in both mice and human data sets (Figure 4C, Chemical drug). When IPA Biological Drug USRs were compared a highly significant correlation again emerged, although only Suhrbier 2 and 4 dpi showed a negative z-score for tocilizumab (Figure 4C, Biological drug; Supplemental Table 6). Why the human data sets all failed to provide a z-score for tocilizumab (Supplemental Table 6) is unclear, given the clear IL-6 signatures (Figure 4A,B). Conceivably, the human lung samples were collected too late, with the best results for tocilizumab achieved when the drug was given early in infection (50). Overall these results argue that K18-hACE2 mice represent a suitable model in which to test biologics and chemotherapeutics for COVID-19.

### Distinct gene expression profiles point to the same dominant cytokine pathways

For human-mouse comparisons, differential gene expression showed poor overlaps (Figure 1B,C), whereas pathway analyses showed highly significant correlations (Figure 3A,B, Figure 4A-C). To dissect this apparent incongruity, the IL-6 receptor signaling network (IL-6R network) was examined in detail. The IL-6 was consistently identified as a USR with positive z-scores (Figure 4A,B), with excessive IL-6 levels also associated with COVID-19 ARDS (49). An IL-6R network, comprising 546 genes, was generated in IPA (Supplemental Table 7A). Each gene (node) was then colored depending on whether it was an up-regulated DEG in one or more mouse data sets (47% of the 546 genes in the network), an up-regulated DEG in 1 or more human datasets (3%), an up-regulated DEG in any human and any mouse data set (27%), or not an up-regulated DEG in any human or mouse data set (23%) (Figure 4D; Supplemental Table 8). The results again illustrated that the overlap in up-regulated DEGs for human-mouse comparisons was relatively low (27%), but also showed that DEGs were spread across the network and sub-networks, with no evidence of species-specific clustering (Fig. 4D). A heat map of log_2_ fold change was generated for the 546 genes in the IL-6R network Figure 4E) and a parallel map was generated to indicate which genes were DEGs (q<0.05) for each data set (Figure 4F). Differences in gene expression patterns were again clearly evident for human and mouse comparisons, even for genes within this dominant pathway (Figure 4E, F).

This analysis was repeated for TNF and IFNg networks, as these were also identified as dominant USRs (Figure 4A, B). Broadly similar results emerged, with many genes only significantly up-regulated in mice, but not humans ((Supplemental Tables 7B, C and Supplemental Figures 5, 6), which might be explained by the higher levels of infection in mice (Figure 2A). However, a number of genes were up-regulated DEGs in humans, but not in mice (Supplemental Figures 5, 6; 18% for TNF and 21% for IFNg); also seen to some extent for the IL-6R network (Figure 4D, 3%). So IL-6R, TNF and IFNg are identified as dominant pathways in both mouse and humans, yet some genes within these pathways were identified as DEGs in humans, but not in mice. This was not primarily because these human DEGs had no orthologues in mice, as this accounted for less than 2% of these genes. The reason these genes were not DEGs in mice was also not because they had significantly higher variance in mice (i.e. higher variance causing lower significance). Instead these genes were not DEGs because their fold-change was significantly lower (Supplemental Figures 7, 8).

Taken together these analyses argue that, although transcriptomic responses may show poor scoDEG overlap between mice and humans, they nevertheless often indicate the activation of common pathways. Said a different way, in mice and humans, dominant pro-inflammatory cytokines/chemokine responses appear to result in the induction of a different, only partially overlapping, set of genes within that pathway.

### Differences in K18-hACE2 backgrounds for Winkler and Suhrbier studies

Perhaps surprising was that the Winkler and Suhrbier datasets did not show a higher levels of concordance (Figure 4E, F; Supplemental Figure 3B). IPA Diseases and Functions analyses for lungs 2 dpi (where mean viral loads were similar, Figure 2A) also showed that cellular infiltrate and immune activation annotations had higher z-scores for the Suhrbier group than for the Winkler group (Table 2). The basis for these differences was unclear given both studies ostensibly used the same inbred K18-hACE2 mice supplied by the Jackson Lab, and both were infected with virus isolates belonging to the original SARS-CoV-2 strain (Table 1).

**Table 2:** IPA ‘Diseases and Functions’ differences between Suhrbier 2 dpi vs. Winkler 2 dpi. DEGs for Suhrbier 2 dpi vs. Winkler 2 dpi. were separately analysed by Diseases and Functions feature of IPA. Diseases and Functions were ranked by the difference in z-scores (Delta z), with the top 30 with the largest differences shown. Any Diseases and Functions that were not significant were nominally given a z-score of 0. Diseases and Functions associated with leukocyte migration are highlighted in grey. On 2 dpi viral loads in lungs were comparable for Suhrbier and Winkler.

| Diseases and Functions | Suhrbier z-score | Winkler z-score | Delta z |
| --- | --- | --- | --- |
| Homing of leukocytes | 5.937 | 0 | 5.937 |
| Recruitment of blood cells | 5.72 | 0 | 5.72 |
| Phagocytosis of leukocytes | 5.499 | 0 | 5.499 |
| Metabolism of reactive oxygen species | 5.33 | 0 | 5.33 |
| Homing of granulocytes | 5.228 | 0 | 5.228 |
| Chemotaxis of granulocytes | 5.228 | 0 | 5.228 |
| Stimulation of cells | 4.991 | 0 | 4.991 |
| Homing of neutrophils | 4.827 | 0 | 4.827 |
| Maturation of blood cells | 4.719 | 0 | 4.719 |
| Migration of lymphatic system cells | 4.606 | 0 | 4.606 |
| Endocytosis by eukaryotic cells | 4.582 | 0 | 4.582 |
| Cell viability of blood cells | 4.518 | 0 | 4.518 |
| Cytotoxicity of cells | 4.451 | 0 | 4.451 |
| Activation of lymphatic system cells | 4.308 | 0 | 4.308 |
| Cell viability of leukocytes | 4.276 | 0 | 4.276 |
| Activation of granulocytes | 4.116 | 0 | 4.116 |
| Differentiation of T lymphocytes | 4.069 | 0 | 4.069 |
| Inhibition of blood cells | 4.057 | 0 | 4.057 |
| Binding of endothelial cells | 4.02 | 0 | 4.02 |
| Response of lymphocytes | 4.016 | 0 | 4.016 |
| Adhesion of immune cells | 6.73 | 2.722 | 4.008 |
| Recruitment of lymphocytes | 3.98 | 0 | 3.98 |
| Adhesion of blood cells | 6.635 | 2.66 | 3.975 |
| Replication of Flaviviridae | 0 | -3.959 | 3.959 |
| Binding of blood cells | 6.836 | 2.888 | 3.948 |
| Homeostasis of leukocytes | 3.865 | 0 | 3.865 |
| Cell movement of lymphatic system cells | 3.849 | 0 | 3.849 |
| Inhibition of cells | 3.83 | 0 | 3.83 |
| Degranulation | 3.826 | 0 | 3.826 |

The K18-hACE2 founder line was created on a mixed C57BL/6J x SJL/J background (51). Perhaps under-appreciated is that C57BL/6J (6J) mice contain a unique loss-of-function deletion of exons 5 to 9 of the *Nicotinamide nucleotide transhydrogenase* (*Nnt* gene), whereas most mouse strains (including SJL/J mice) encode a full length *Nnt* gene (52). An exact-match k-mer method targeting exon 2 and 9 of the *Nnt* gene was used to interrogate the RNA-Seq data from the Suhrbier and Winkler studies. Most mice from the Suhrbier data sets had no exon 9 reads (Figure 5A), consistent with a dominant 6J background, with these mice maintained in-house as heterozygotes by repeated backcrossing onto 6J mice. In contrast, all of the K18-hACE2 mice from the Winkler study had exon 9 reads, arguing that these mice had an intact *Nnt* gene and that this line had not been extensively backcrossed onto 6J.

**Figure 5:**
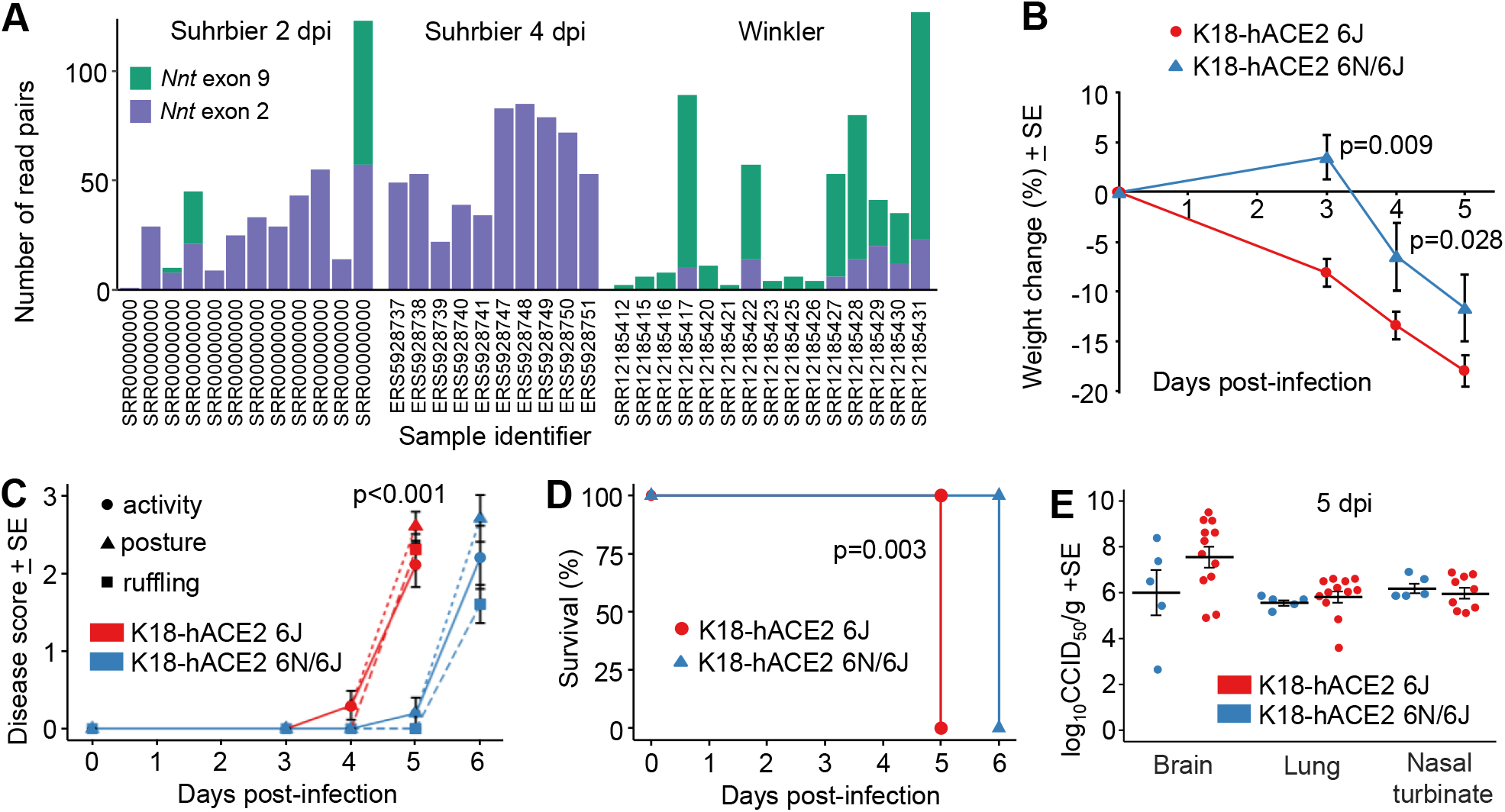
Genetic background of K18-hACE2 mice and disease progression. **(A)** For each sample of the Suhrbier and Winkler datasets, the number of read pairs originating from exon 9 and exon 2 of the *Nnt* gene are shown as green and purple bars, respectively. **(B)** Change in body weight over five days following SARS-CoV-2 infection is shown as a percentage of starting body weight for K18-hACE2 6J and K18-hACE2 6J/6N mice; p-values indicate significant difference between means at 3 and 4 dpi (Kruskal-Wallis tests, n=5 per group). **(C)** Three disease score parameters (activity, posture and fur ruffling) over six days following SARS-CoV-2 infection is shown for K18-hACE2 6J and K18-hACE2 6J/6N mice. Statistics for 5 dpi for all 3 parameters by Kolmogorov-Smirnov tests (n=5 per group). **(D)** Kaplan-Meier curves showing survival for K18-hACE2 6J and K18-hACE2 6J/6N mice following SARS-CoV-2 infection. Significance by log rank test (n=5 per group). **(E)** Log_10_CCID_50_/g in brain, lung and nasal turbinate for K18-hACE2 6J and K18-hACE2 6J/6N mice 5 dpi with SARS-CoV-2. 6J data was derived from 2 independent experiments. Differences between K18-hACE2 6J and K18-hACE2 6J/6N mice for any tissue were not significance.

To explore further the influence of genetic background, we undertook a single back-cross of our K18-hACE2 mice (6J background, *Nnt^-/-^*) with C57BL/6N (6N) mice (*Nnt*^+/+^), and compared SARS-CoV-2 infection in (i) K18-hACE2 mice on 6J background (K18-hACE2 6J, *Nnt*^-/-^) with (ii) K18-hACE2 mice on a mixed 6J x 6N background (K18-hACE2 6J/6N, *Nnt^+/-^*). Weight loss, disease scores and survival were all significantly worse in K18-hACE2 6J mice than in the K18-hACE2 6N/6J mice (Figure 5B-D). The genetic background can thus influence SARS-CoV-2 induced inflammatory immunopathology in the K18-hACE2 model (Table 2); an observation consistent with a previous report that showed such a mixed background can also significantly affect an arthritic inflammatory immunopathology (52). The *Nnt* gene may, at least in part, be responsible as NNT controls mitochondrial reactive oxygen species via the glutathione and thioredoxin pathways, and thus generally exerts an anti-inflammatory influence (52). No significant differences in tissues titres were observed (Figure 5E), consistent with previous studies which also showed no detectable influence of such a mixed background on antiviral activity against an alphavirus (52).

In summary, the differences seen for the Winkler vs. Suhrbier datasets can be explained by the differences in the genetic backgrounds of the K18-hACE2 mice.

### The mACE2-hACE2 mouse model

A criticism of the K18-hACE2 mouse model has been that hACE2 expression is driven by the keratin 18 promoter, which *inter alia* results in a fulminant brain infection that is associated with mortality (30). An alternative, less severe, non-lethal model of SARS-CoV-2 infection involves use of transgenic mice where expression of hACE2 is driven by the mouse ACE2 promoter (mACE2-hACE2 mice) (34). We have independently generated this mouse model (Supplemental Figure 9A,B) and show that lung titers in mACE2-hACE2 mice were lower than those seen in K18-hACE2 mice on day 2 post infection (≈2 logs) (Figure 6A). Weight loss was also less prominent reaching ≈ 7% by day 8, with K18-hACE2 mice approaching 20% by day 5 (Supplemental Figures 9C). Importantly, although nasal turbinates are infected (Supplemental Figures 9D) there were no detectable brain infections (Figure 6A). Lung histology shows characteristic loss of alveolar spaces, cellular infiltrates, smooth muscle hypertrophy/hyperplasia, and bronchial sloughing (Supplemental Figure 9E), although this lung pathology is less severe than that seen in K18-hACE2 mice (28). Our mACE2-hACE2 mice thus behave similarly to those described by Bao et al. 2020 (34).

**Figure. 6.**
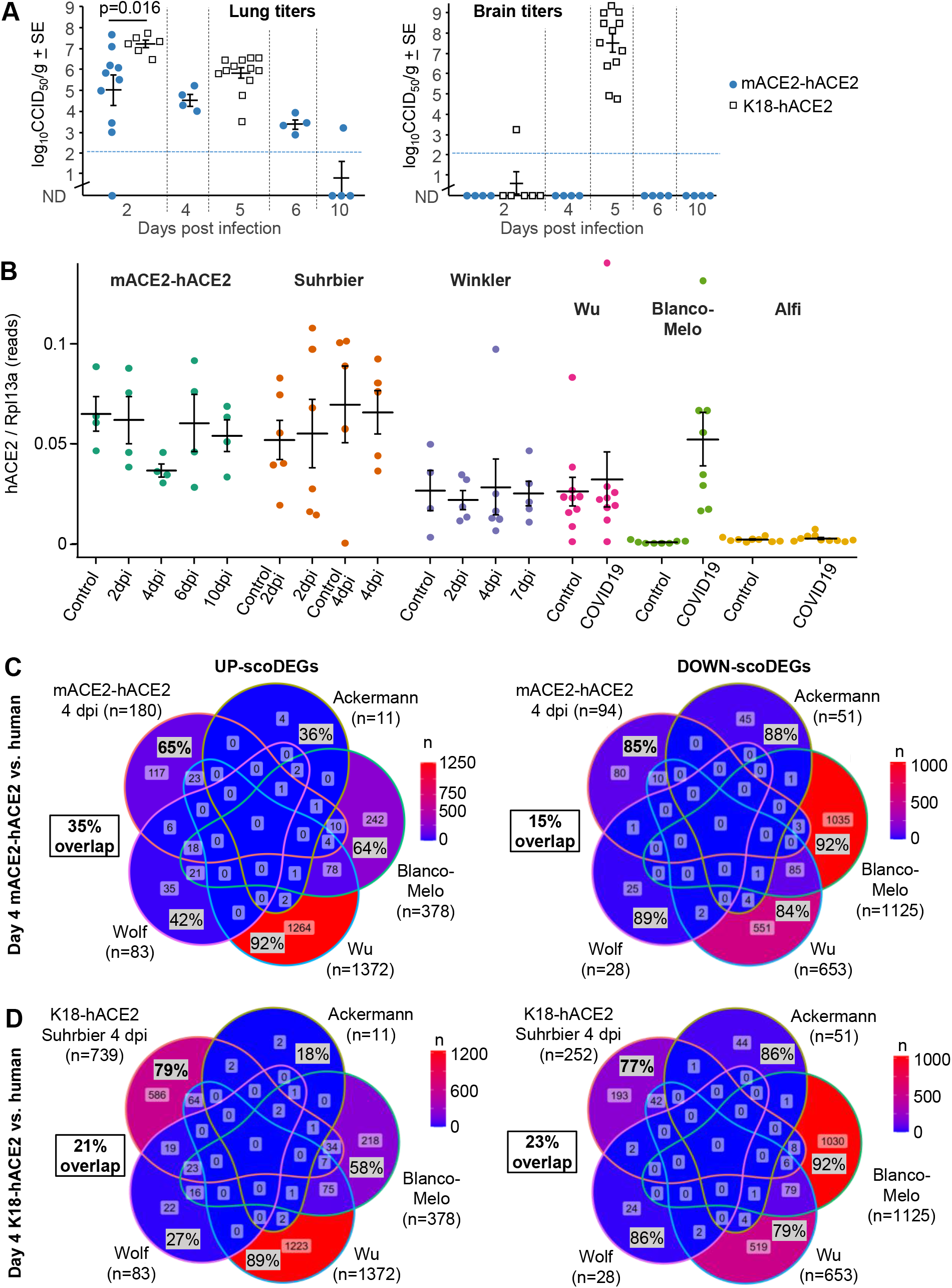
The mACE2-hACE2 mouse model. **(A)** Lung and brain viral tissue titers at the indicated days post infection. All infected K18-hACE2 mice reach ethically defined endpoints for euthanasia (weight loss >20%) by day 5. None of the mACE-hACE2 mice reach ethically defined endpoints for euthanasia. Mean lung titers on day 2 were 2.2 log_10_CCID_50_ lower in mACE2-hACE2 mice compared to (Suhrbier) K18-hACE2 mice (p=0.016, Kolmogorov-Smirnov test). (**B)** hACE2 reads normalized to Rpl13a reads for all mACE2-hACE2, K18-hACE2 and human lung samples. Cross-bars represent group means + standard error. **(C)** Venn-diagrams show overlap in up- and down-regulated scoDEGs between mACE2-hACE2 4 dpi mice and the four human groups. Percentages within the Venn diagram (grey boxes) show the percentage of scoDEGs exclusive to that group (i.e. a scoDEG in that group, but in no other group). The boxed overlap percentages represent the percentage of 4 dpi mouse scoDEGs that are also scoDEGs in one or more human studies (e.g. 180-117/180 × 100 ≈ 35% for up-regulated scoDEGS and 94-80/94 ×100 ≈15% for down-regulated scoDEGs). **(D)** As for C showing overlaps in up- and down-regulated scoDEGs between K18-hACE2 Suhrbier 4 dpi and the four human groups. Percentages as in C.

### RNA-Seq of infected mACE2-hACE2 mouse lungs

RNA-Seq analyses of SARS-CoV-2 infected mACE2-hACE2 mice was undertaken for days 0, 2, 4, 6 and 10 post infection (Table 1). Expression levels of hACE2 mRNA in this model were not significantly different from those seen in K18-hACE2 mice (Figure 6B, mACE2-hACE2 and Suhrbier). The reduced viral loads 2 dpi in mACE2-hACE2 mouse lungs (Figure 6A) and lower pathogenicity was thus not due to overall lower levels of receptor expression in lungs (Figure 6B). hACE2 levels in the other groups (Figure 6A) may not be strictly comparable as *inter alia* the Winkler study sequenced total RNA, the Wu study sequenced ribo-depleted RNA (rather than poly adenylated RNA), the Blanco-Melo study had low read depth (Supplemental Figure 1), and the Alfi data was derived from organoids (Table 1).

The RNA-Seq analysis of infected mACE2-hACE2 mouse lungs provided a series of DEGs, with day 4 post infection providing the highest number of DEGs (Supplemental Table S9). The scoDEG overlap for mACE2-hACE2 (4 dpi) vs. human studies was 35% for up-regulated scoDEGs and 15% for down-regulated scoDEGs (Figure 6C). The same comparisons for K18-hACE2 Suhrbier (4 dpi) vs. human studies provided 21% and 23% (Figure 6D), and for K18-hACE2 Winkler (4 dpi) vs. human studies gave 33% and 27%, respectively Supplemental Figure 10A). Thus, despite the lower vial loads (Figure 6A), overall scoDEG overlap for mACE2-hACE2 mice vs. human groups, was not consistently better than those seen for K18-hACE2 mice vs. human groups. PC1/PC2 plots support the contention that concordance between mACE2-hACE2 and human groups shows no substantial improvement over K18-hACE2 vs. human groups (Supplemental Figure 10B).

The overlap for up-regulated DEGs for 4 dpi mACE2-hACE2 vs. 4 dpi K18-hACE2 was high, 76% (Supplemental Figure 10C), with both strains on a 6J background. The concordance for IPA Cytokine USRs for this comparison was also highly significant (Supplemental Figure 10D). These data argue that the promoter that drives hACE2 expression does not play a major role in determining the nature of the inflammatory responses. The lower overlap in down-regulated DEGs (30%, Supplemental Figure 10C) may be associated with the differences in viral loads (Figure 6A), but may also reflect different viral tropisms, given that hACE2 is driven by distinct promoters.

### Pathway comparisons between infected mACE2-hACE2 and human lungs

The number of DEGs obtained from mACE2-hACE2 infected lungs was substantially lower than that obtained from K18-hACE2 infected lungs (Supplemental Table 9 vs. Supplemental Table 1). Nevertheless, when analyzed by IPA, inflammatory pathways again (as for K18-hACE2 mice, Figure 4A-C) showed a high level of concordance with human studies (Figure 7A-C). Chemical Drug annotations again included dexamethasone (Figure 7C, Chemical Drug). No annotations for Biological Drug were identified, perhaps reflecting the mild nature of SARS-CoV-2 infection and the lower number of DEGs for this model.

**Figure 7:**
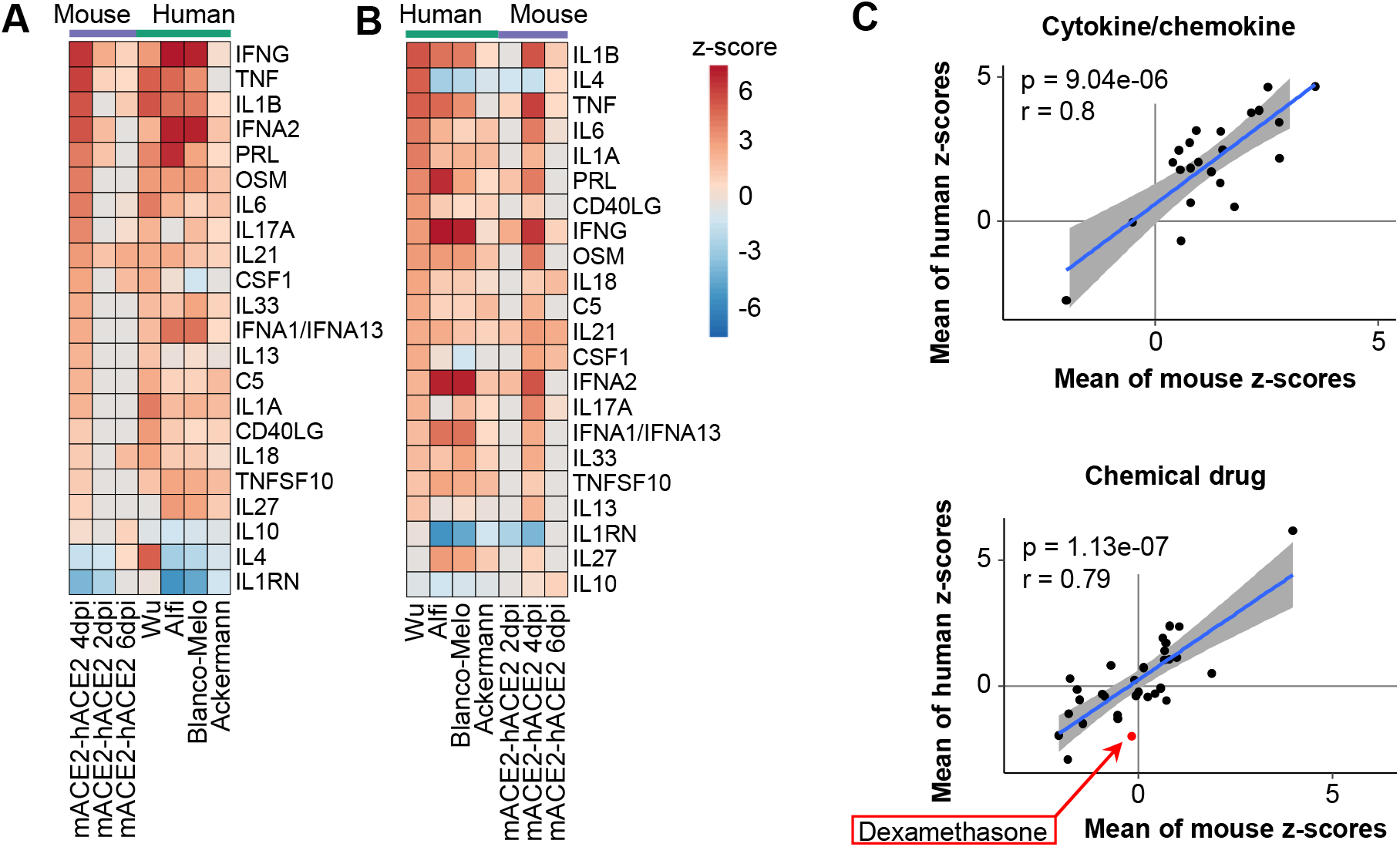
Cytokine/chemokine and drug USR concordances between mACE2-hACE2 mice and humans. **(A)** Heatmap comparing mACE2-hACE2 mouse groups with human groups for IPA cytokine/chemokine USRs ranked by activation z-score in mACE2-hACE2 4 dpi. **(B)** Heatmap comparing groups as in A, except ranked according to z-score in Wu, as a representative of human groups. **(C)** Pearson correlation of mean mouse vs. mean human z-scores for significantly enriched cytokine/chemokine USRs (n=22), and chemical drug USRs (n=30). Blue line shows linear regression with 95% confidence intervals (grey).

## Discussion

We show herein that two mouse models of SARS-CoV-2 infection, one severe (K18-hACE2) and one mild (mACE2-hACE2), both recapitulated human gene expression quite poorly. SARS-CoV-2 infection in mouse and human lung tissues induced transcriptomic reponses that overlapped by 21-35% for up-regulated scoDEGs and 15-27% for down-regulated scoDEGs (Figure 1B, 6C; Supplementary Fig10A)). In contrast, species concordance for inflammatory pathways and immune-related signatures was highly significant (Figs. 2B, 3A,B, 4A-D and 7). Mice and humans would thus appear to regulate individual genes within such pathways somewhat differently, but overall many of the same pathways are activated. As dominant pathways in immunity and inflammation are the target of most COVID-19 interventions, these mouse models can be viewed as providing representative and pertinent models for pre-clinical assessments of new interventions.

The K18 promoter in K18-hACE2 mice drives receptor expression in cells that ordinarily would not express ACE2 (19). The mACE2-hACE2 mice might be viewed as more physiologically relevant, in that receptor expression is restricted to cells where the mACE2 promoter is active. Brain infection is therefore ostensibly lost in mACE2-hACE2 mice presumably because neuroepithelial cells (53) do not express hACE2. Perhaps surprisingly, the lower levels of viral infection in the lungs in mACE2-hACE2 mice was not associated with lower overall hACE2 mRNA expression levels (Figure 6B). One might speculate that in K18-hACE2 mice, certain lung cells aberrantly express ACE2 and become infected (54, 55)} leading to higher viral loads. Either way, the poor overlap in DEGs for K18-hACE2 groups and human studies was not demonstrably due to hACE2 expression being driven by the non-physiological K18 promoter, as the poor overlap was largely retained for mACE2-hACE2 mice. Furthermore, the inflammatory responses seen in infected mACE2-hACE2 and K18-hACE2 lungs were very similar (Supplemental Figure 10D), arguing that the promoter is also not a major factor in determining the nature of the inflammatory responses.

There are clearly a number of limitations for this kind of analysis. Unavoidable is the issue of single copy orthologues, which comprised 63-77% of genes identified by RNA-Seq in lung tissues. This issue is less of a problem for pathway analyses when using programs such as IPA that accept both human and mouse gene nomenclature. The different sources of tissues and the different technologies used to generate gene expression data (Table 1) likely add to non-biological variability, although this was perhaps mitigated herein by combining multiple human and mouse studies. The large differences in viral loads between some groups (Figure 2A) would appear to play a role in the poor concordance in gene expression profiles, particularly for human groups and down-regulated genes. However, analyses of K18-hACE2 (high viral load) and mACE2-hACE (lower viral loads) argued that the difference in viral loads was not a major player in the poor overlap in up-regulated orthoDEGs for mouse vs. human groups.

In summary, the analyses herein argue that overlap in orthoDEG expression in the lung tissues of hACE2-transgenic mice and humans after SARS-CoV-2 infection is generally poor. In contrast, the concordance in immune and inflammation pathways was high, arguing that the transgenic mouse models provide relevant and pertinent models in which to evaluate new interventions for SARS-CoV-2 and COVID-19.

## Methods

### Ethics statement and regulatory compliance

All mouse work was conducted in accordance with the “Australian code for the care and use of animals for scientific purposes” as defined by the National Health and Medical Research Council of Australia. Mouse work was approved by the QIMR Berghofer Medical Research Institute animal ethics committee (P3600, A2003-607). For intrapulmonary inoculations via the intranasal route, mice were anesthetized using isoflurane. Mice were euthanized using CO2. All infectious SARS-CoV-2 work was conducted in a dedicated suite in a biosafety level-3 (PC3) facility at the QIMR Berghofer MRI (Australian Department of Agriculture, Water and the Environment certification Q2326 and Office of the Gene Technology Regulator certification 3445).

### Virus preparation

The SARS-CoV-2 isolate (hCoV-19/Australia/QLD02/2020) was kindly provided by Dr Alyssa Pyke and Fredrick Moore (Queensland Health Forensic & Scientific Services, Queensland Department of Health, Brisbane, Australia). The virus sequence is deposited at GISAID (https://www.gisaid.org/) (29). Virus stocks were prepared in Vero E6 cells as described (29) and were checked for mycoplasma as described (56). The fetal calf serum used for propagation of cells and virus was checked for endotoxin contamination as described (57).

### K18-hACE2 mice

K18-hACE2+/- mice were purchased from Jackson laboratories and were maintained in-house as heterozygotes by backcrossing to C57BL/6J mice (27, 28). Mice were typed as described (29) using hACE2 Primers: Forward: 5’-CTT GGT GAT ATG TGG GGT AGA -3’; Reverse: 5’-CGC TTC ATC TCC CAC CAC TT -3’ (recommended by NIOBIOHN, Osaka, Japan).

### mACE2-hACE2 mice

These mice were generated by Monash Genome Modification Platform (MGMP), Monash University and are freely available through Phenomics Australia (MGMP code ET26). Briefly, the mouse BAC clone RP23-152J15 was obtained from BACPAC Genomics and was used to generate a mACE2 promoter subclone. hACE2 cDNA with a polyA tail was cloned into mACE2 promoter subclone. The hACE2 (ENSG00000130234) sequence was codon optimized for mouse expression and was ordered as a synthetic cDNA with homology arms (GeneArt). The transgenic construct contained the mACE2 promoter and hACE2 followed by a poly A (Supplemental Figure 9A). A maxiprep was then digested with Asc1 and the extracted 10,062 fragment (2.5 ng/ml) microinjected into the pronucleus of C57BL/6J zygotes at the pronuclei stage. Injected zygotes were transferred into the uterus of pseudo pregnant F1 females.

Mice were genotyped using the following primers 5’-TCC GGC TGA ACG ACA ACT CC - 3’, 5’-TAT GTT TCA GGT TCA GGG GGA GG -3’. Cycling conditions were: 1 cycle at 94°C for 3 mins; 35 cycles of 94°C for 30 secs, 60°C for 30 secs, and 72°C for 1 mins; and 1 cycle of 72°C for 10 min followed by cooling to 4°C. Fragments were run on a gel with a 374 bp band indicating the presence of the transgene (Supplemental Figure 9B). The mACE2-hACE2 mouse line was maintained in-house as heterozygotes by backcrossing onto C57BL/6J mice.

### Mouse infections

Mice were infected intrapulmonary via the nasal route with 5×10^4^ CCID_50_ of virus in 50 μl medium while under light anesthesia; 3% isoflurane (Piramal Enterprises Ltd., Andhra Pradesh, India) delivered using The Stinger Rodent Anesthesia System (Advanced Anaesthesia Specialists/Darvall, Gladesville, NSW, Australia).

Mice were scored daily on a scale of 0-3 according to posture, activity, fur ruffling, and fever. For all criteria, the normal condition was designated as 0. For posture, hunching only while at rest was designated as 1, moderate hunching with some impairment of normal movement was designated as 2, and severe hunching with difficulty in maintaining upright posture was designated as 3. For activity, a mild to moderate decrease was designated as 1, stationary unless stimulated was designated as 2, and reluctant to move even if stimulated was designated as 3. For fur ruffling and fever, mild to moderate fur ruffling was designated as 1, severe ruffling was designated as 2, and shivering was designated as 3. Any animal reaching a level of 3 in any single criterion was euthanized, and any animal reaching a level of 2 in two or more criteria was euthanized.

Body weight was measured daily. Viral titrations were performed at 5 days post-infection with a CCID_50_ assay using Vero E6 cells and serial dilution of supernatants from homogenized tissues as described previously (29).

### Gene expression analysis

Suitable human and mouse COVID-19 transcriptome datasets were identified by searching the National Centre for Biotechnology Information Sequence Read Archive (NCBI-SRA) via the BigQuery platform using the command: ‘SELECT distinct m.bioproject FROM nih-sra-datastore.sra.metadata as m, UNNEST (m.attributes) as a WHERE (m.organism=‘Homo sapiens’ OR m.organism=‘Mus musculus’) AND assay_type=‘RNA-Seq’ AND ((a.v LIKE ‘SARS%2’) OR (a.v LIKE ‘COVID%’))’. The search was also extended to include micro-array data and non-publicly available data by searching the NCBI-PubMed database using search terms: ‘COVID’ and ‘SARS’. Microarray data relating to Ackermann et al. (2020) were accessed via the Vivli Centre for Global Clinical Research Data.

Raw sequence data for Winkler, Wu, Blanco-Melo and Alfi groups (see Table 1) were accessed from SRA using fasterq-dump from SRA-toolkit. Quality control of fastq files was performed using FastQC v0.11.9 (58). Adapter sequences were identified using BBmerge from the BBmap package v38.90 (59), and FastQC. Reads were trimmed to remove adapter content, size-selected to remove reads less than 36nt in length, and quality-filtered to remove reads with less than a Q20 Phred score within a sliding-window tetramer, using Trimmomatic v0.36 (60). Processed reads were aligned to either the GRCm39 vM26 or GRCh38 v37 reference genome for mouse and human datasets, respectively, using STAR aligner v2.7.1a (61). Prior to alignment, each reference genome was augmented to include the NC_045512 SARS-Cov-2 Wuhan-Hu-1 viral genome. The number of reads mapping to SARS-CoV-2 was calculated using Samtools v1.9 (62). For paired-end datasets, only primary proper pairs were counted. Host gene expression was calculated using RSEM v1.3.1 (63) and differential expression was calculated using Bioconductor v3.13 (64) and EdgeR v3.34.0 (65) in R v4.1.0 (66). Genes with read coverage of less than 2 counts per million were excluded from all further analyses. Mouse-human orthologues were extracted from the Ensembl database using BiomaRt v2.48.2 (67) in R. Following read-alignment, it was noted that the Blanco-Melo et al. data had very low sequencing depth in COVID-19 infected samples. Therefore, gene expression data were obtained from Supplementary Table 6 of Blanco-Melo et al. (38).

Approximately 20% of human-mouse orthologues have gene IDs that differ between species. Prior to performing any cross-species comparison of scoDEGs, all mouse IDs were changed to human. Overlap between groups for DEGs and scoDEGs was calculated in R and plotted using Eulerr v6.1.0 (68) in R. Pearson’s correlation of log_2_ fold-changes using the union of scoDEGs for each pairwise combination of groups was calculated in R. In instances where data was missing for a scoDEG in one group (due to failing CPM >2 threshold for RNA-Seq data or being absent from the immune gene panel for Ackermann data), the scoDEG was excluded from that pairwise correlation. The proportion of up- and down-regulated DEGs and scoDEGs shared between groups was calculated in R and plotted using ggVennDiagram v1.1.4 (69) in R. Mean mouse log_2_ fold-change and mean human log_2_ fold-change were compared by Pearson’s correlation using scoDEGs that were significant in at least one human group and/or at least one mouse group. In instances where data was missing for a scoDEG in a particular group, log_2_ fold-change of that scoDEG was made zero for that group.

### Reciprocal gene set enrichment analysis

For each group, a log_2_ fold-change ranked gene list was produced using DESeq2 (70) with default settings. Also for each group, orthoDEG sets were filtered to retain only the top 50% of orthoDEGs when ranked according to absolute log_2_ fold change. For all ranked gene lists and filtered orthoDEG sets, gene IDs of mouse-human orthologues were standardized by substituting mouse IDs for their human equivalent where gene IDs differed between species. A Gene Set Enrichment Analysis using GSEA v4.1.0 (40) with 100 permutations and the ‘no_collapse’ setting was used to test for enrichment of filtered orthoDEG sets within ranked gene lists.

### Immune-SigDB gene set enrichment analysis

For each group, log_2_ fold change ranked gene lists were produced as described above, except that no standardization was performed on gene IDs of mouse-human orthologues. The Immune-SigDB v7.4 (15A) gene set collection comprising 5219 immune-related gene sets was obtained from the Molecular Signatures Database (71). A gene set enrichment analysis was performed as described above to test for enrichment of Immune-SigDB gene sets within ranked gene lists. Any gene set not found to be significantly enriched in a particular ranked gene list was given a NES of zero for that group.

### Pathway analysis

Pathway analysis was performed using Ingenuity Pathway Analysis (IPA) v65367011 (Qiagen) with default settings. Data were plotted using pheatmap v1.0.12 (72) and ggplot2 v3.3.3 (73) in R. Gene networks were constructed using USR output and the My Pathways tool in IPA. For each of the IL-6R, TNF, and IFNg USRs, a list of ‘Molecules in dataset’ was obtained for each group. ‘Molecules in dataset’ from each group were then concatenated to create a single molecule list related to each of the three USRs of interest. Each molecule list was used as input to the My Pathways tool. Starting with each USR, the ‘Build/Grow’ function was used to identify direct and indirect downstream relationships between that USR and any of the molecules in its respective molecule list. The following parameters were set as follows: ‘Data Sources’ was set to all; ‘Confidence Level’ was set to ‘Experimentally observed’; ‘Species’ was set to human and mouse; all ‘Tissues and Cell Lines’ were selected except for those relating to cancer; all ‘Relationship Types’ were selected. To identify targets of transcription factors, a second round of ‘Build/Grow’ was performed in same manner as the first except that only direct relationships were allowed, and relationships had to include the transcription factors identified in the first round of ‘Build/Grow’. Each network was then exported in tabular format and plotted using Cytoscape v3.8.2 (74).

### Nnt genotyping

Mouse RNA-Seq data were interrogated for the presence of exon two and nine of the nicotinamide nucleotide transhydrogenase (*Nnt*) gene as described (52) using Repair and BBduk from the BBmap package v38.90. Reads containing at least one 31-mer exactly matching either exon were counted as belonging to that exon. If one member of a read pair matched an exon, the other member was also counted as a match.

### Statistics

Statistics were performed using IBM SPSS Statistics for Windows, version 19.0. For gene expression data a Pearson’s correlation test was used when the number of observations was high (>1000) in accordance with the central limit theorem, or residuals were normally distributed (Shapiro-Wilk test and quantile-quantile plot); otherwise the Spearman’s rank test was used.

For mouse data (weight change, disease scores, virus titers) the non-parametric Kolmogorov–Smirnov or Kruskal-Wallis tests were used as the data was not normally distributed (i.e. differences in variance was >4, skewness < −2, and/or kurtosis was >2). Survival was compared between two groups using a log rank test.

## Author contributions

CRB, TD and DJR undertook the bioinformatic analyses. DJR, TTL, KY and BT undertook the mouse work. GH provided advice on statistical analyses and approaches and undertook statistical analyses. AS and DJR secured the funding. AS wrote the manuscript with input from all the authors.

## Acknowledgements

We thank Drs Alyssa Pyke and Fredrick Moore (Queensland Health, Brisbane) for providing SARS-CoV-2. For generation of the mACE2-hACE2 mice, the authors acknowledge the facilities, and the scientific and technical assistance of the Monash Genome Modification Platform (MGMP), Monash University. MGMP is supported by Phenomics Australia (PA). PA is supported by the Australian Government through the National Collaborative Research Infrastructure Strategy (NCRIS) program. From QIMR Berghofer MRI, we thank Dr Itaru Anraku for managing the PC3 (BSL3) facility and animal house staff for mouse breeding and agistment, and Dr V Lutzky for proof reading. This publication is based in part on research using data made available by Vivli, Inc. Vivli has not contributed to, or approved, and is not in any way responsible for, the contents of this publication.

