## Supplemental Figures for "Mouse models of COVID-19 recapitulate inflammatory pathways rather than gene expression"

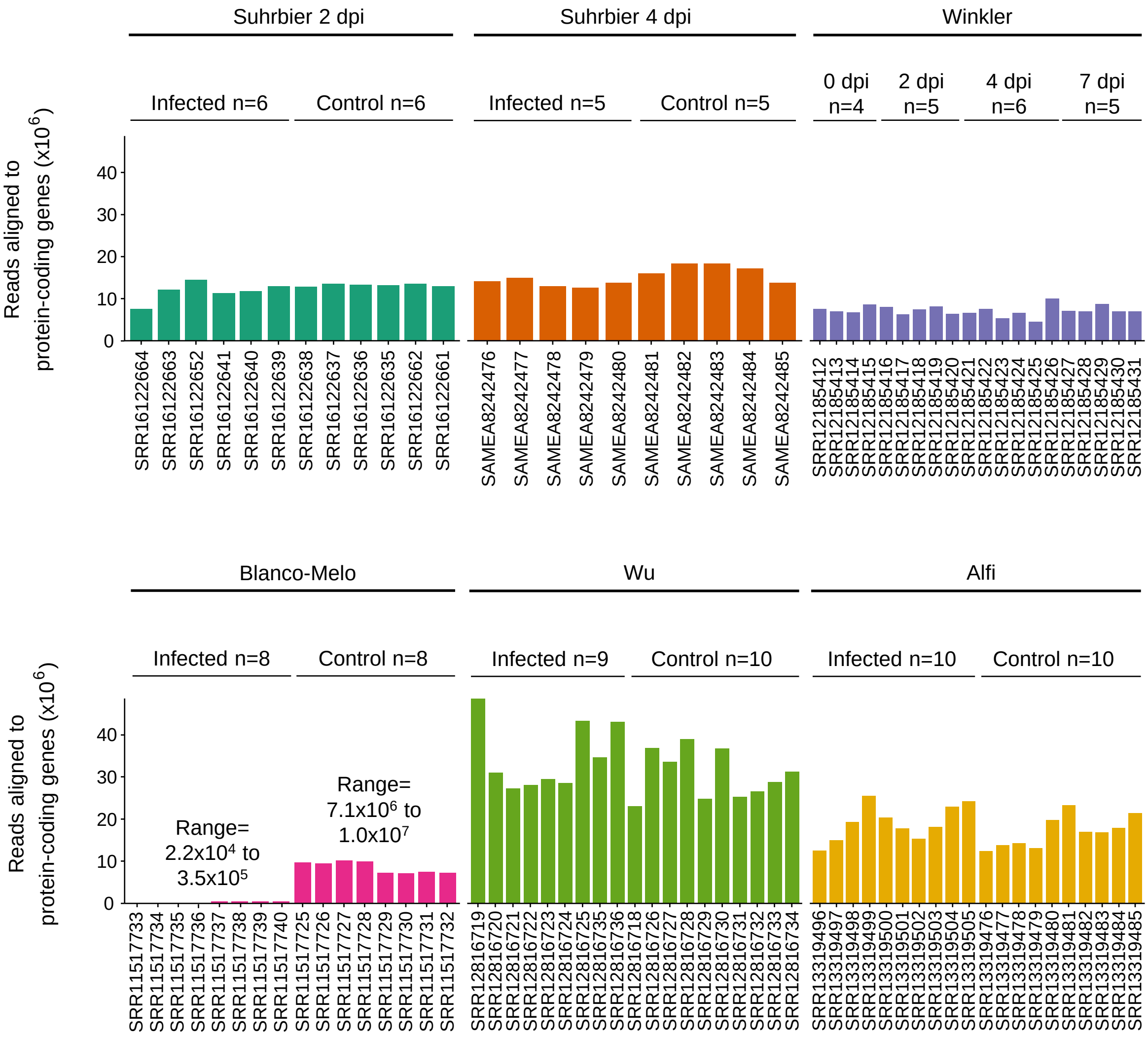


**Supplemental Figure 1. Number of RNA-Seq reads aligned to protein-coding genes.** For each sample, reads were aligned to either the mouse GRCm39 M26 or human GRCh38 v37 reference genome using STAR. Reads aligning to protein-coding genes were counted using RSEM. The total number of reads aligned to protein coding genes are shown for each sample. Due to low coverage in Blanco-Melo infected samples, read data were not re-analysed for this dataset. Instead, differential expression results were obtained from the original publication (Blanco-Melo et al., 2020).

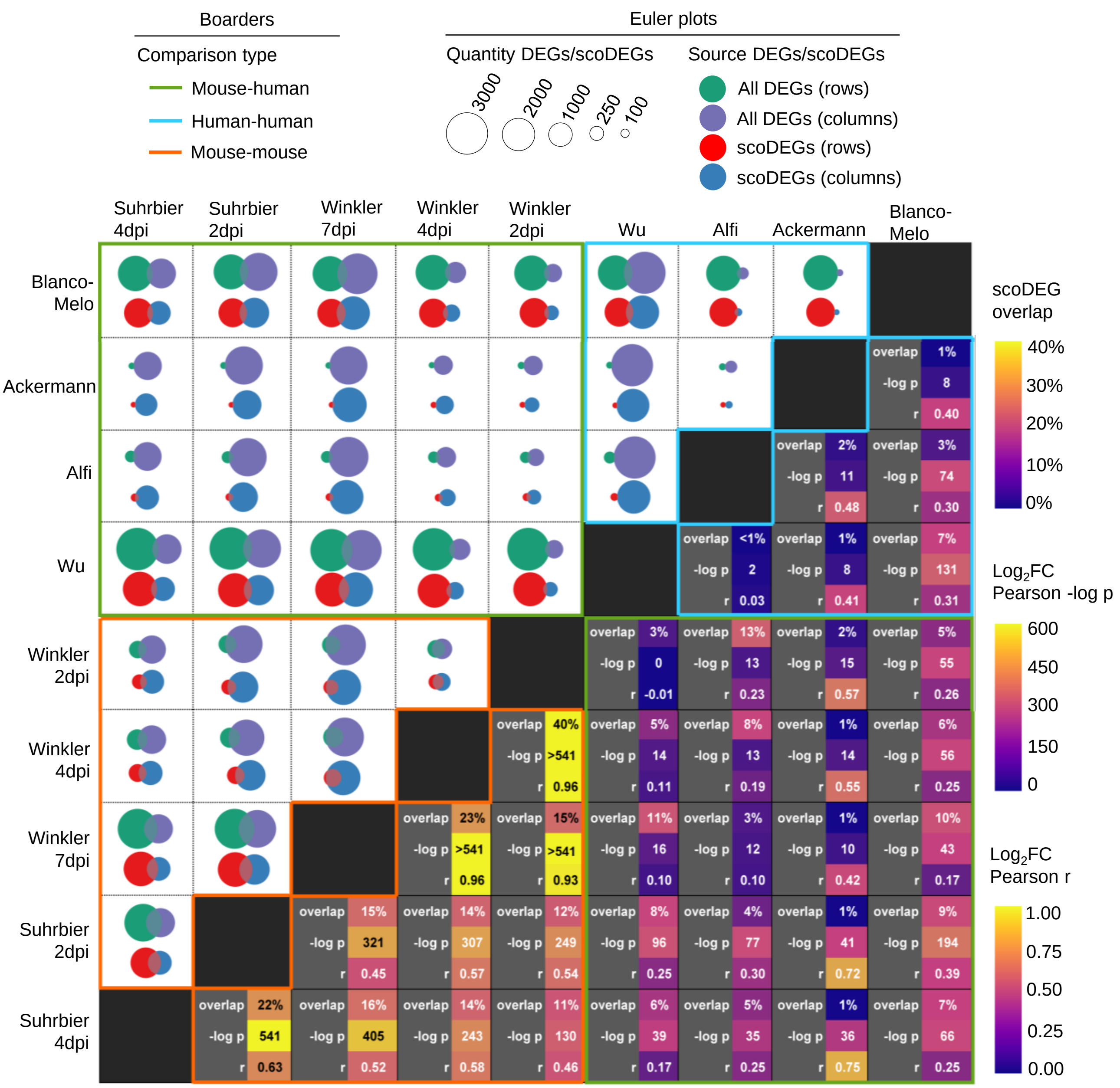


**Supplemental Figure 2. Pair-wise comparisons between groups of differential gene expression. Upper-left** Euler diagrams show the amount of overlap between groups regarding DEGs (green and purple circles) and scoDEGs (red and blue circles) for all possible group-wise combinations. Green and red circles relate to row names, while purple and blue circles relate to column names. Size of circles indicates the number of DEGs/scoDEGs, as produced by EdgeR analysis or, in the case of Ackermann and Blanco-Melo, as obtained from the authors. **Lower-right** Each cell contains information pertaining to the group-wise comparison indicated by the row and column names. Overlap - for each pair-wise comparison between groups the number of scoDEGs that were common to both groups is shown as a percentage of the total number of scoDEGs in the comparison. –log p and r - for each pair-wise comparison, gene expression was compared using the union of scoDEGs for those groups (i.e. single-copy orthologues that were differentially expressed in one or both groups, and that were present in the gene lists for both groups). Pearson correlations were then performed using the $\log_2$ fold-changes (log2FC) of those single-copy orthologues to provide –log p and r values. Ackerman provides high r values as this analysis only evaluated expression of 249 inflammation genes (see Table 1). Cells are colored using a scales on the right.  For upper left and lower right, colored boarders indicate whether comparisons are mouse-human (green), human-human (blue), or mouse-mouse (orange).

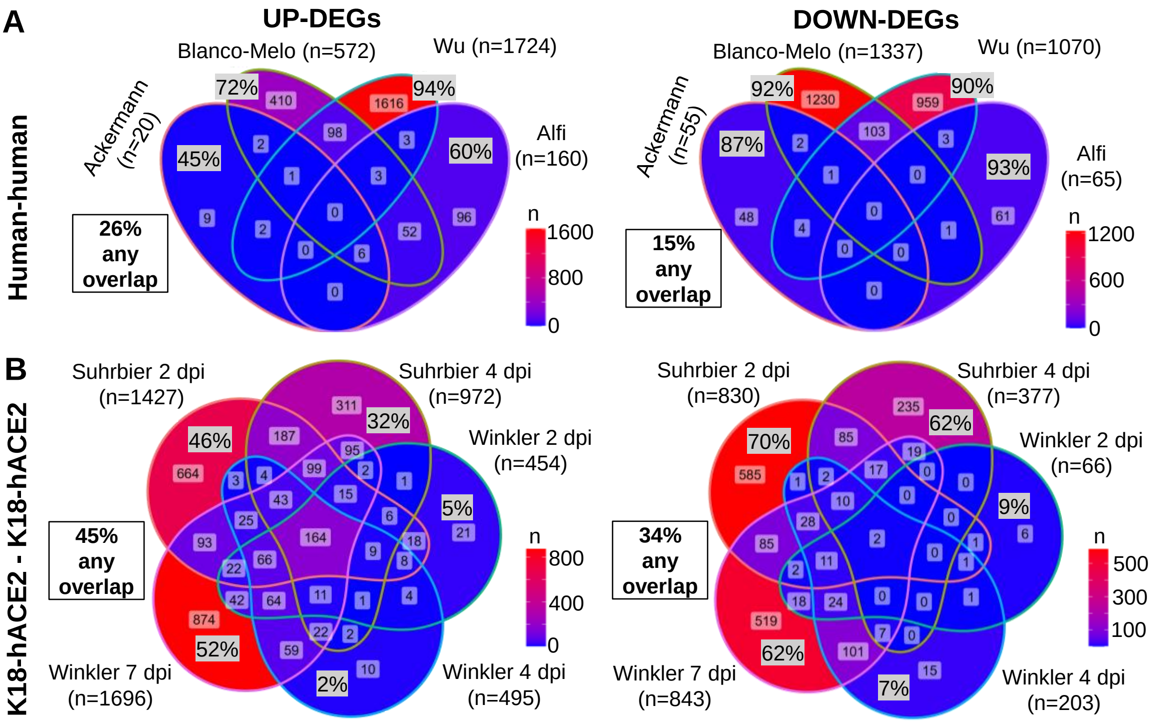


**Supplemental Figure 3. (A)** All human groups were compared for overlap of up- and down-regulated DEGs. 'n' refers to the number of DEGs for each group. Within each segment of each Venn diagram the percentage of DEGs exclusive to that group (i.e. a DEG in that group but no other group) is provided as a percentage of the total number of DEGs in that group (e.g. 9/20 x 100 = 45%). The boxed percentages (any overlap) refer to the percent of all DEGs in the Venn that are shared by at least 2 groups. **(B)** As for A, except comparing DEGs between all K18-hACE2 mouse groups.

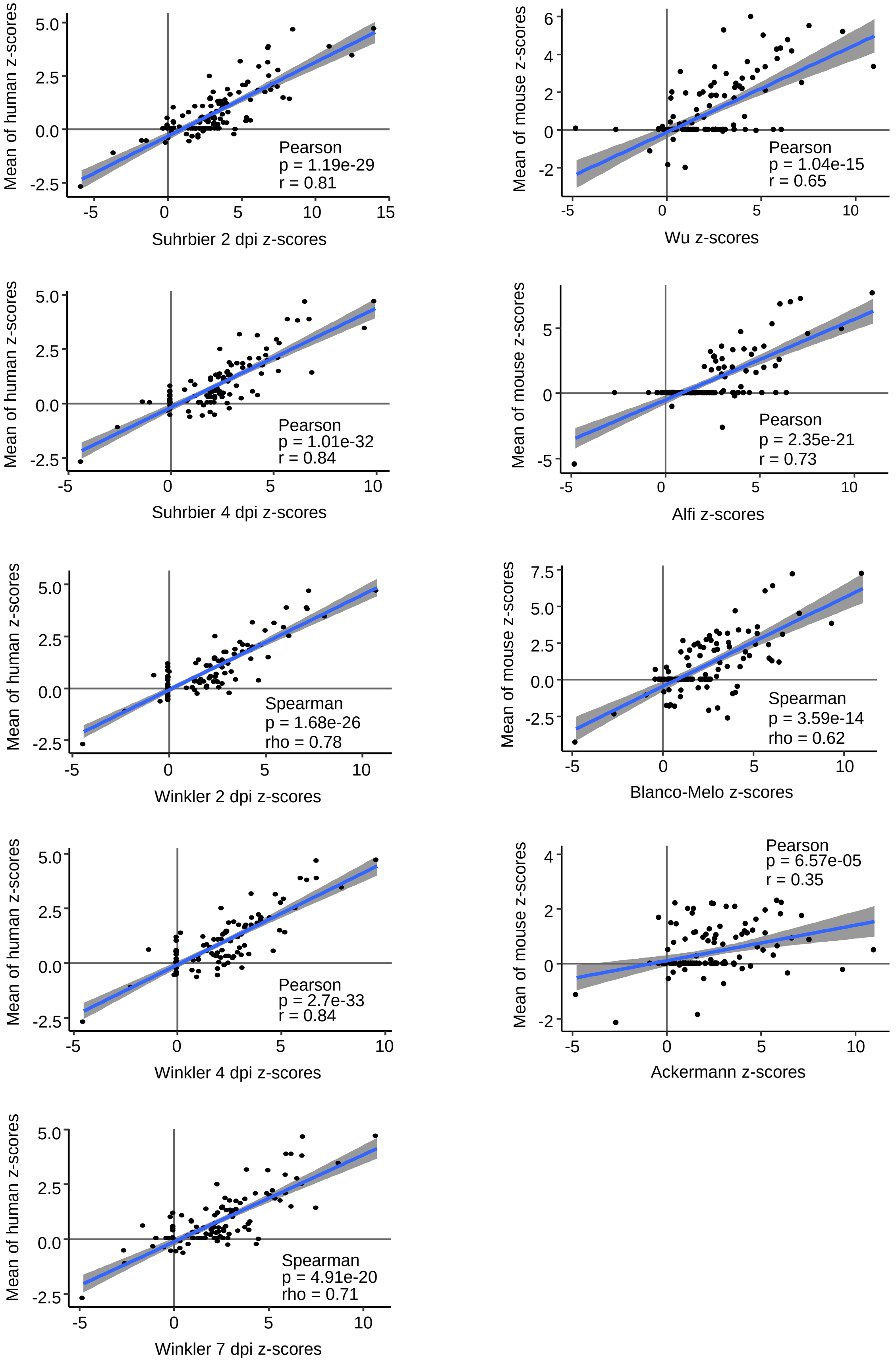


**Supplemental Figure 4. Correlation of group z-score vs. mean z-score for comparing cytokine/chemokine USR activation between species.** Activation z-scores for each group are plotted on x-axes (left column = mouse groups, right column = human groups). Mean z-scores for each species are plotted on y axes (left column = mean of all human groups, right column = mean of all mouse groups).

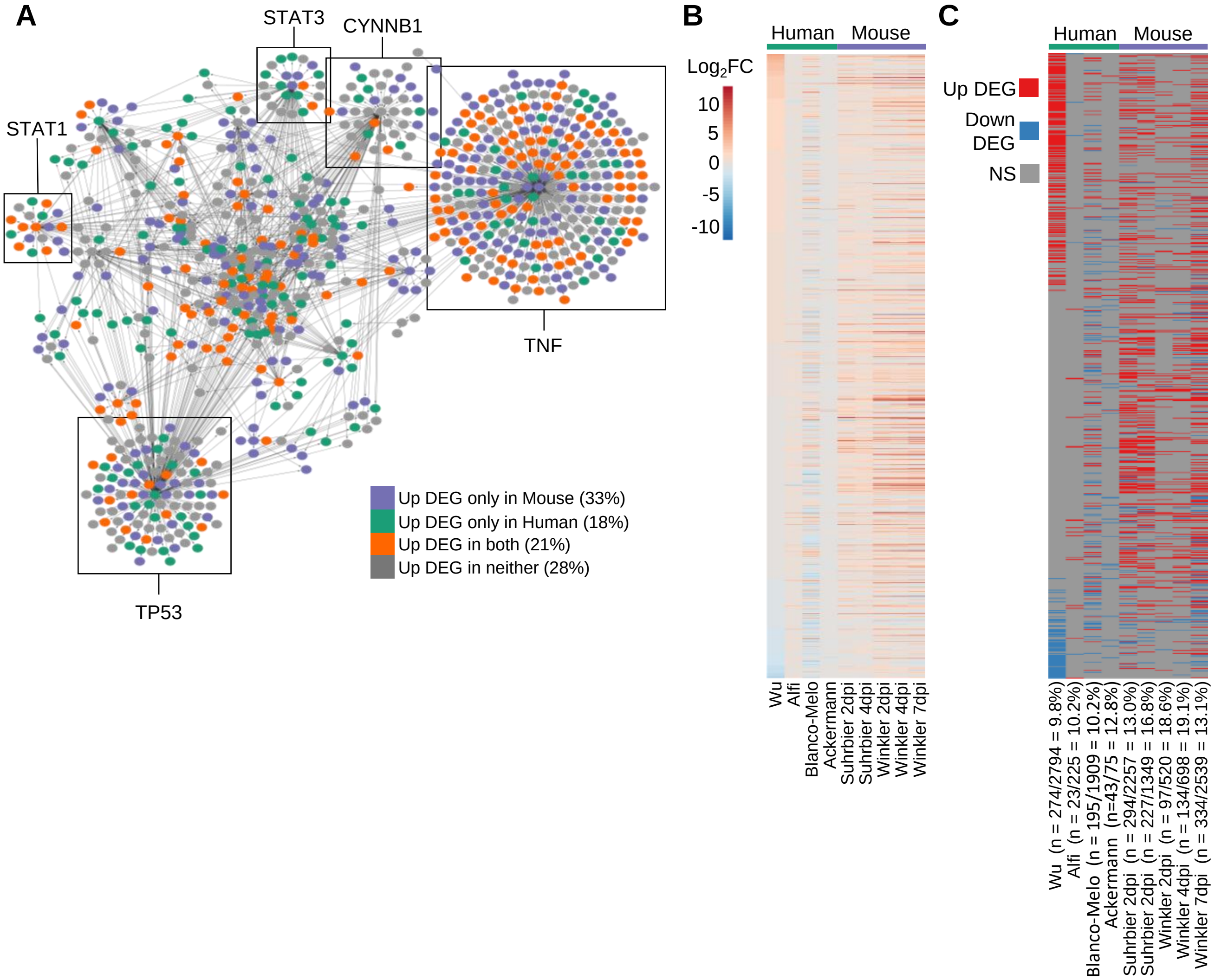


**Supplemental Figure 5. Differential expression of 1000 genes associated with TNF signaling. (A)** Regulatory network for TNF signaling was constructed in the following manner: DEGs from each group were used as input for a separate IPA Core Analysis. The 'Upstream Regulators' output was used to identify genes associated with TNF signaling. Results from all groups were concatenated into a single list of 1000 genes. This list was used to interrogate DEG lists from each group in order to identify which TNF-associated genes were up-regulated in each group. Node colour indicates whether a gene was up-regulated in mouse only (>1 mouse group, and no human), human only (>1 human group, and no mouse), both (> 1 mouse and > 1 human group), or none. Large sub-networks are labeled according to their hub node. **(B)** Heatmap comparing groups according to $\log_2$ fold-change ($\log_2$FC) of 1000 genes associated with TNF signaling. Genes are ordered according to $\log_2$FC in Wu. **(C)** Heatmap comparing groups according to differential expression of 1000 genes associated with TNF signaling. Genes are ordered as in B. Cells are coloured according to whether the gene was significantly up-regulated (red), significantly downregulated (blue), or not significant (NS, grey). The number of TNF genes that were significantly differentially expressed is shown for each group as a percentage of the total number of DEGs for that group (n).

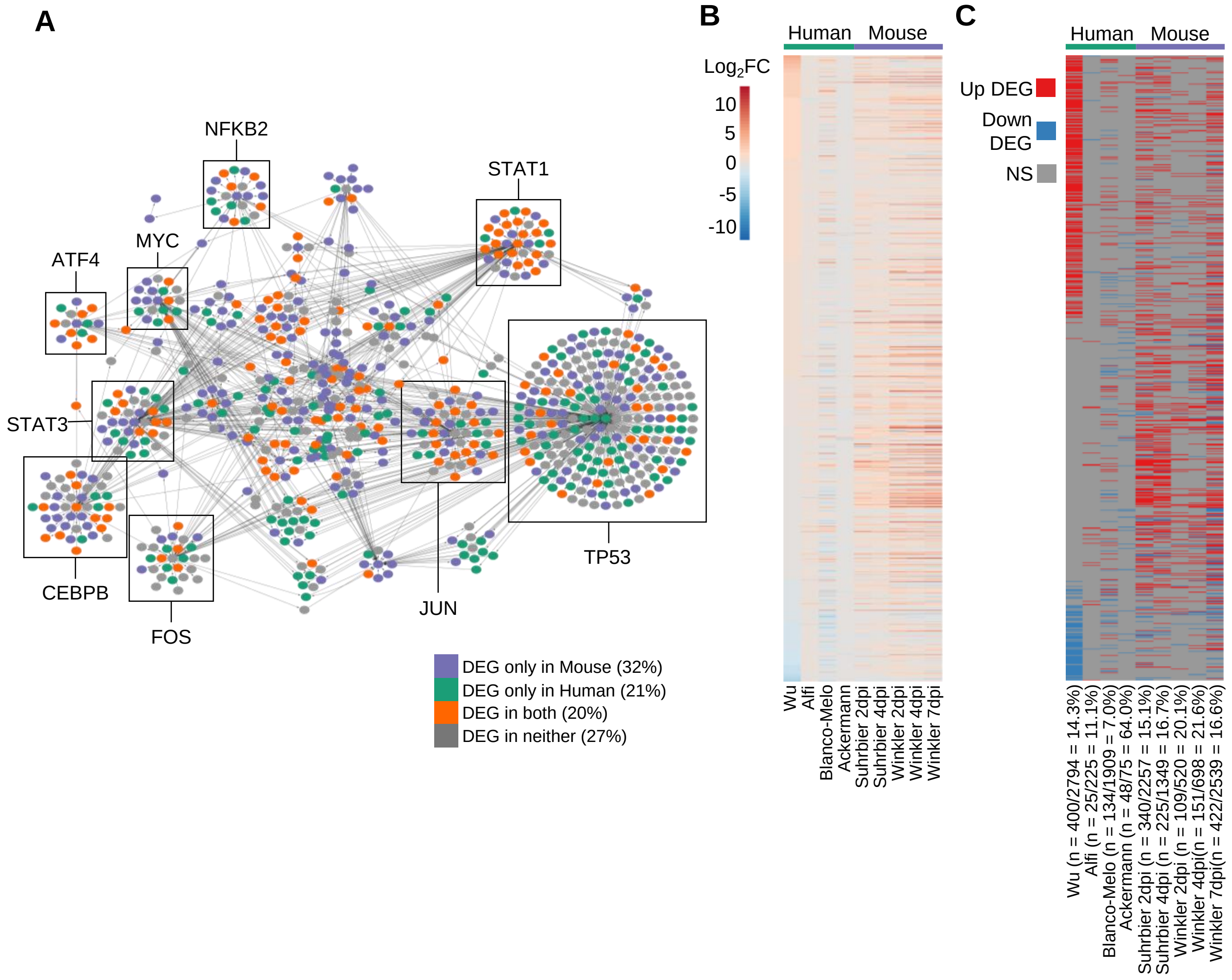


**Supplemental figure 6. Differential expression of 862 genes associated with IFNg signaling. (A)** Regulatory network for IFNg signaling was constructed in the following manner: DEGs from each group were used as input for a separate IPA Core Analysis. The 'Upstream Regulators' output was used to identify genes associated with IFNg signaling. Results from all groups were concatenated into a single list of 862 genes. This list was used to interrogate DEG lists from each group in order to identify which IFNg-associated genes were up-regulated in each group. Node colour indicates whether a gene was up-regulated in mouse only (>1 mouse group, and no human), human only (>1 human group, and no mouse), both (> 1 mouse and > 1 human group), or none. Large sub-networks are labeled according to their hub node. **(B)** Heatmap comparing groups according to $\log_2$ fold-change ($\log_2$FC) of 862 genes associated with IFNg signaling. Genes are ordered according to $\log_2$FC in Wu. **(C)** Heatmap comparing groups according to differential expression of 862 genes associated with IFNg signaling. Genes are ordered as in B. Cells are coloured according to whether the gene was significantly up-regulated (red), significantly downregulated (blue), or not significant (NS, grey). The number of IFNg genes that were significantly differentially expressed is shown for each group as a percentage of the total number of DEGs for that group (n).

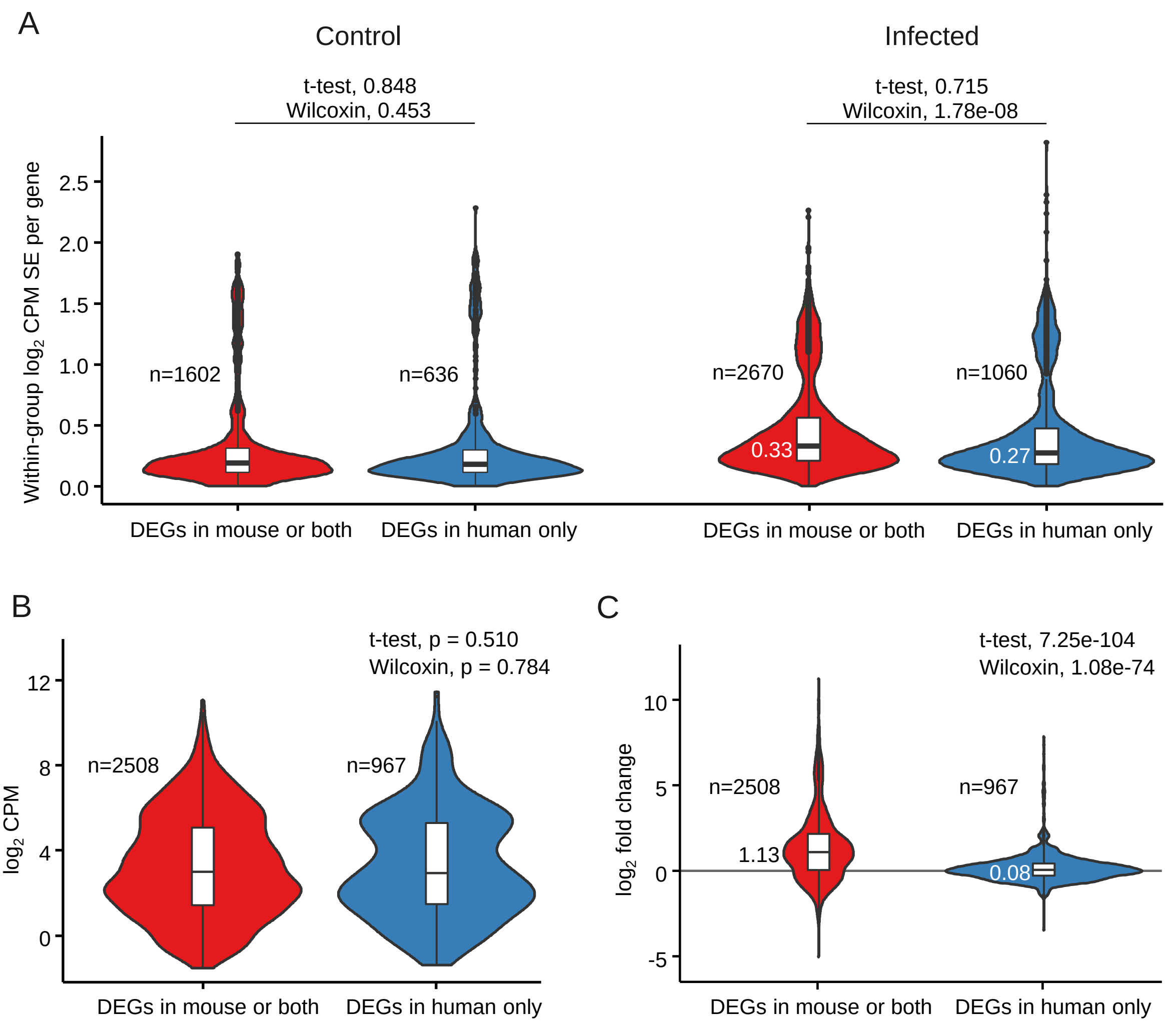


**Supplemental Figure 7. Potential causes for lack of significant differential expression of TNF-related genes in mice. (A)** To examine the effect of natural variability on differential expression, mouse orthologues that were DEGs in mouse or both mouse and human ($\geq$1 mouse groups and $\geq$0 human groups) were compared to those that were DEGs in human only (0 mouse groups and $\geq$1 human groups). This was done separately for Infected and Control. For each gene, the standard error of $\log_2$ TMM-normalised counts per-million ($\log_2$ CPM SE) was calculated. There are five Infected mouse groups (Suhrbier 2 dpi, Suhrbier 4 dpi, Winkler 2 dpi, Winkler 4 dpi, and Winkler 7 dpi), and three Control mouse groups (Suhrbier 2 dpi, Suhrbier 4 dpi, and Winkler 0 dpi). Therefore ‘n’ represents the number of genes multiplied by the number of groups in that treatment (e.g. n=1500 refers to 500 genes * 3 Control groups). Numbers shown in white refer to median values. For comparisons, medians and means were tested for significant difference between ‘groups’ using a Wilcoxin test and a Welch’s t-test respectively. **(B)** To examine the effect of read depth on differential expression, mouse orthologues that were DEGs in mouse or both mouse and human were compared to those that were DEGs in human only, in terms of their $\log_2$ CPM across all eight mouse groups (five Infected and three Control). Therefore ‘n’ represents the number of genes multiplied by eight). Tests for significant difference were performed as in A. **(C)** Mouse orthologues that were DEGs in mouse or both mouse and human were compared to those that were DEGs in human only, in terms of $\log_2$ fold-change. Numbers shown in white refer to median values. Tests for significant difference were performed as in A.

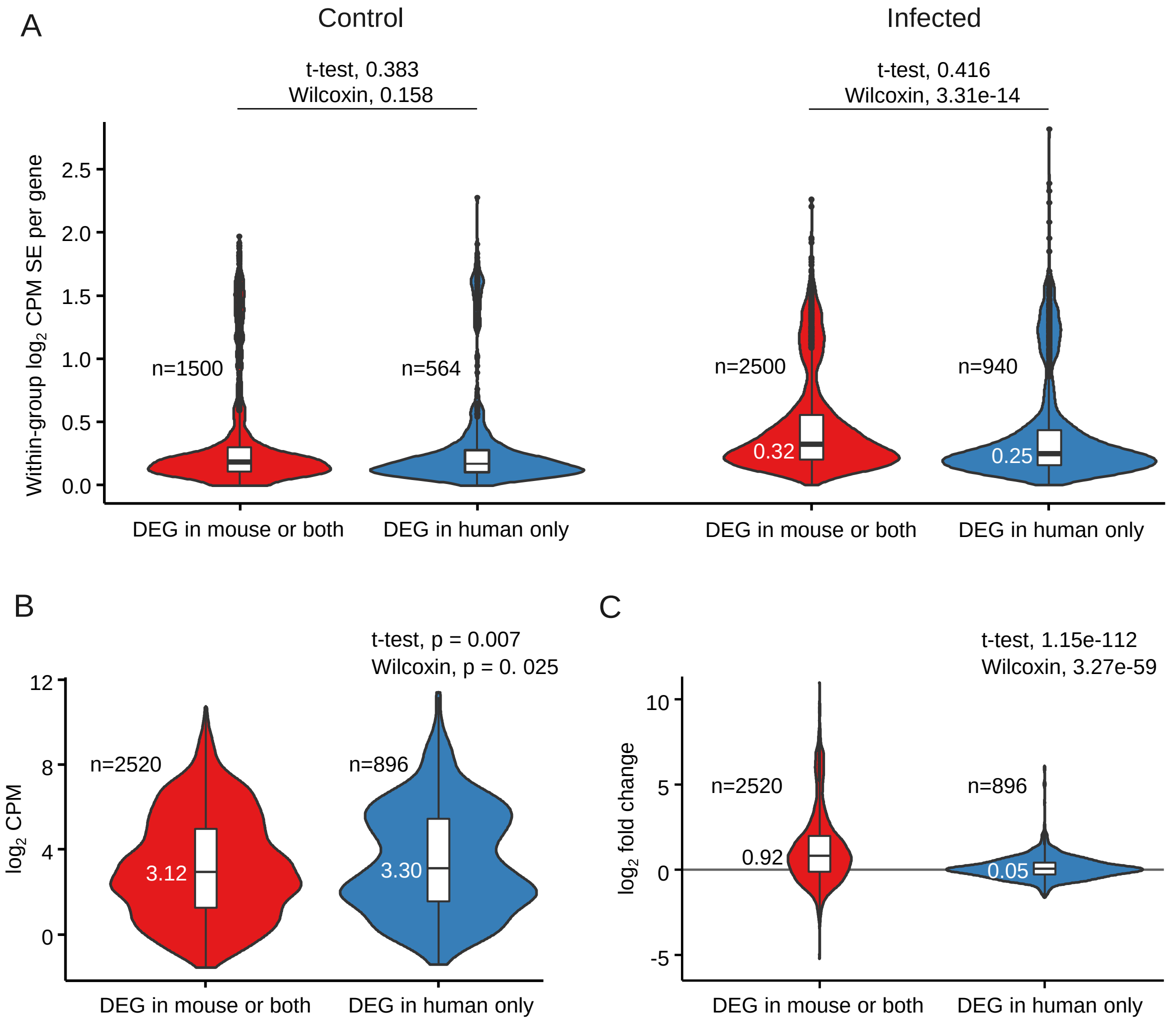


**Supplemental Figure 8. Potential causes for lack of significant differential expression of IFNg-related genes in mice. (A)** To examine the effect of natural variability on differential expression, mouse orthologues that were DEGs in mouse or both mouse and human ($\geq$1 mouse groups and $\geq$0 human groups) were compared to those that were DEGs in human only (0 mouse groups and $\geq$1 human groups). This was done separately for Infected and Control. For each gene, the standard error of $\log_2$ TMM-normalised counts per-million ($\log_2$ CPM SE) was calculated. There are five Infected mouse groups (Suhrbier 2 dpi, Suhrbier 4 dpi, Winkler 2 dpi, Winkler 4 dpi, and Winkler 7 dpi), and three Control mouse groups (Suhrbier 2 dpi, Suhrbier 4 dpi, and Winkler 0 dpi). Therefore ‘n’ represents the number of genes multiplied by the number of groups in that treatment (e.g. n=1602 refers to 534 genes * 3 Control groups). Numbers shown in white refer to median values. For comparisons, medians and means were tested for significant difference between ‘groups’ using a Wilcoxin test and a Welch’s t-test respectively. **(B)** To examine the effect of read depth on differential expression, mouse orthologues that were DEGs in mouse or both mouse and human were compared to those that were DEGs in human only, in terms of their $\log_2$ CPM across all eight mouse groups (five Infected and three Control). Therefore ‘n’ represents the number of genes multiplied by eight). Tests for significant difference were performed as in A. **(C)** Mouse orthologues that were DEGs in mouse or both mouse and human were compared to those that were DEGs in human only, in terms of $\log_2$ fold-change. Numbers shown in white refer to median values. Tests for significant difference were performed as in A.

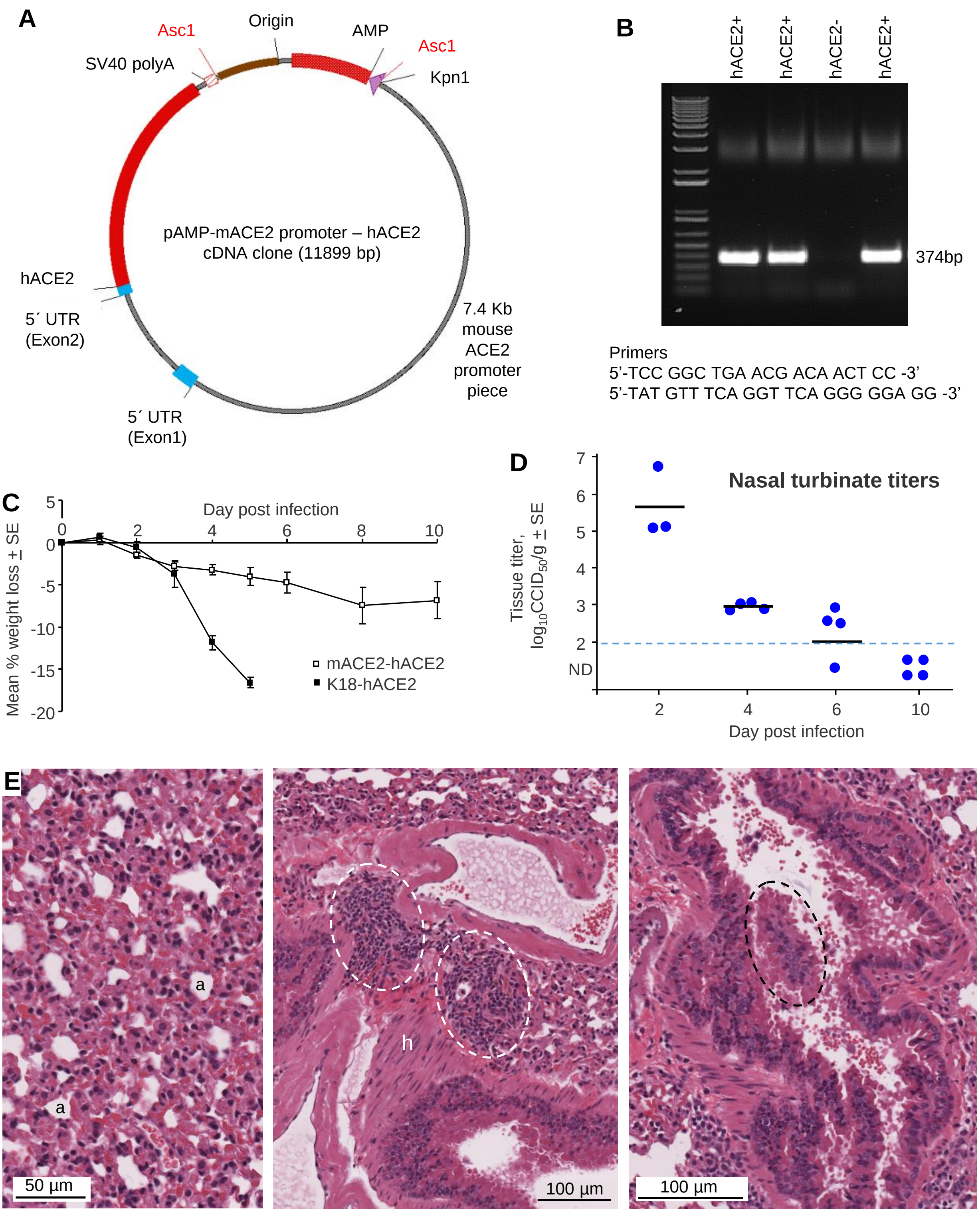


**Supplemental Figure 9. mACE2-hACE2 mice. (A)** The transgenic construct used for generation of mACE2-hACE2 mice containing the mACE2 promoter and hACE2 followed by a poly A. **(B)** Genotyping transgenic mice, a 374 bp PCR fragment indicates the presence of hACE2. **(C)** mACE2-hACE2 mice (n=16 on day 0) were weighed at the indicated times, with 4 mice euthanized on days 2, 4, 6 and 10. K18-hACE2 mice were infected with the same dose of SARS-CoV-$2_{QLD02}$ (n=8) and were all euthanized on day 5. **(D)** Nasal turbinate tissue titers on the indicated days post infection. Limit of detection ≈2 $log_{10}CCID_{50}$/g (ND – ND detected). **(E)** Lung H&E 6 dpi showing loss of alveolar spaces (a - remaining spaces) (left), cellular infiltrates (white dashed ovals), smooth muscle hypertrophy/hyperplasia (h), and bronchial sloughing (black dashed oval).

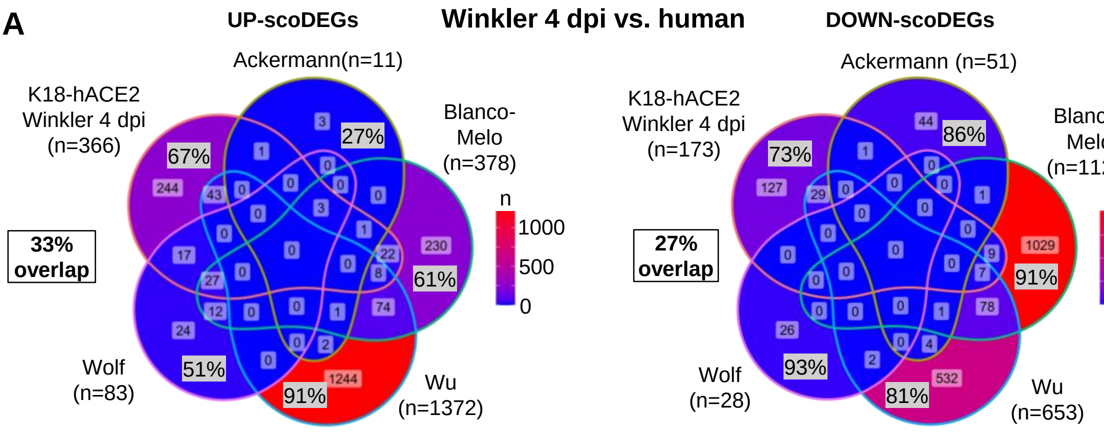


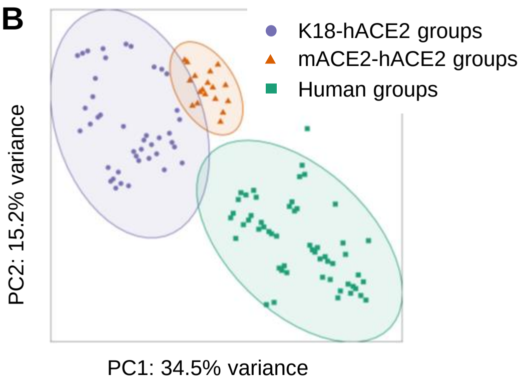


**Supplemental Figure 10. scoDEG and DEG overlaps between groups. (A)** Venn-diagrams show overlap in up- and down-regulated scoDEGs between Winkler 4 dpi and four human groups. Boxed overlap percentages represent the overlap in scoDEGs between K18-hACE2 Winkler 4 dpi and any human study. **(B)** Principal Component Analysis plots including all RNA-Seq data sets; ovals represent 95% confidence interval.

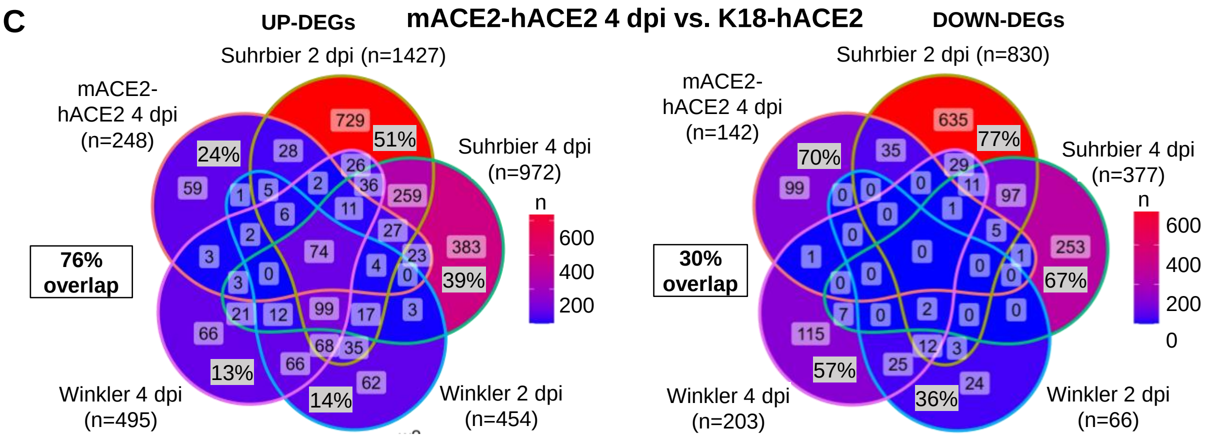


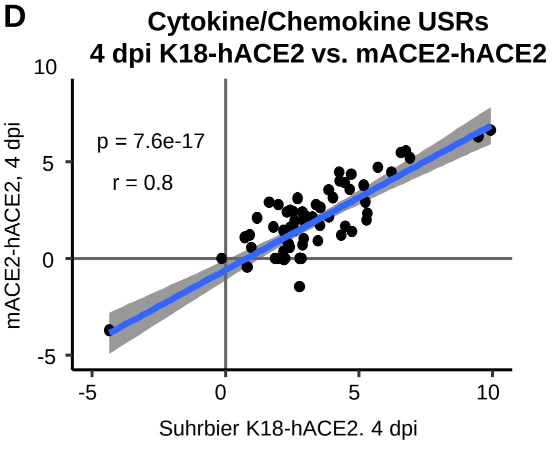


**Supplemental Figure 10. (C)** Venn-diagrams show overlap in up- and down-regulated DEGs between mACE2-hACE2 4 dpi and K18-hACE2 Suhrbier and Winkler groups on 2 and 4 dpi. Boxed overlap percentages represent the overlap in mACE2-hACE2 4 dpi DEGs and DEGs in any of the indicated K18-hACE2 groups. (**D**) Cytokine USRs (n=70) from IPA plotted as in Fig. 4C, but comparing 4 dpi from K18-hACE2 and mACE2-hACE2.
